# A genetic screen to identify factors affected by undecaprenyl phosphate recycling uncovers novel connections to morphogenesis in *Escherichia coli*

**DOI:** 10.1101/2020.07.28.225441

**Authors:** Matthew A. Jorgenson, Joseph C. Bryant

## Abstract

Undecaprenyl phosphate (Und-P) is an essential lipid carrier that ferries cell wall intermediates across the cytoplasmic membrane in bacteria. Und-P is generated by dephosphorylating undecaprenyl diphosphate (Und-PP). In *Escherichia coli*, BacA, PgpB, YbjG, and LpxT dephosphorylate Und-PP and are conditionally essential. To identify vulnerabilities that arise when Und-P metabolism is defective, we developed a genetic screen for synthetic mutations in combination with Δ*ybjG* Δ*lpxT* Δ*bacA*. The screen uncovered system-wide connections, including novel connections to cell division, DNA replication and repair, signal transduction, and glutathione metabolism. Further analysis revealed several new morphogenes; loss of one of these, *qseC*, caused cells to enlarge and lyse. QseC is the sensor kinase component of the QseBC two-component system. In the absence of QseC, the QseB response regulator is overactivated by PmrB cross-phosphorylation. Here, we show that deleting *qseB* completely reverses the shape defect of Δ*qseC* cells, as does overexpressing *rprA* (a small RNA). Surprisingly, deleting *pmrB* only partially suppressed *qseC*-related shape defects. Thus, QseB is activated by multiple factors in the absence of QseC and functions ascribed to QseBC may be related to cell wall defects. Altogether, our findings provide a framework for identifying new determinants of cell integrity that could be targeted in future therapies.

## Introduction

Most glycan layers that surround and protect bacteria are assembled from monomers linked to a polyisoprene lipid carrier (Manat *et al.*, 2014); for the majority of bacteria, this is undecaprenyl phosphate (Und-P), a 55-carbon isoprene commonly referred to as bactoprenol. Und-P is essential by virtue of its requirement to synthesize a stress-bearing structure known as the peptidoglycan (PG) sacculus (Kato *et al.*, 1999, Bouhss *et al.*, 2008, Manat *et al.*, 2014), a net-like macromolecule that surrounds the cytoplasmic membrane and confers upon bacteria their tell-tale shape (Vollmer *et al.*, 2008, Typas *et al.*, 2011). Specifically, nucleotide activated precursors UDP-*N*-acetylglucosamine and UDP-*N*-acetylmuramyl pentapeptide are synthesized in the cytoplasm and are assembled on Und-P to form the PG precursor lipid II (Egan *et al.*, 2020). Lipid II is then translocated across the cytoplasmic membrane by a flippase (Sham *et al.*, 2014, Meeske *et al.*, 2015) and polymerized into glycan chains by PG glycosyltransferases and cross-linked by PG transpeptidases (Egan *et al.*, 2020). The cellular pool of Und-P is generated from the dephosphorylation of Und-PP (undecaprenyl pyrophosphate) to Und-P by integral pyrophosphatases. The BacA and PAP2 protein families produce the monophosphate linkage from Und-PP synthesized in the cytoplasm via the non-mevalonate pathway (Hunter, 2007, Manat *et al.*, 2014) or from Und-PP released on the outer side of the cytoplasmic membrane during glycan polymerization (El Ghachi *et al.*, 2004, El Ghachi *et al.*, 2005). In the Gram-negative bacterium *Escherichia coli*, BacA provides the majority of phosphatase activity (El Ghachi *et al.*, 2004), with smaller contributions provided by the PAP2 enzymes PgpB, YbjG, and LpxT (formerly YeiU) (El Ghachi *et al.*, 2005).

Multiple lines of evidence demonstrate that ongoing Und-PP dephosphorylation is essential to maintain a sufficient supply of Und-P acceptor molecules to properly expand the PG sacculus without fatal breaches. For example, depleting BacA from an *E*. *coli* mutant lacking YbjG and PgpB causes cells to lyse (El Ghachi *et al.*, 2005). Similarly, depleting BcrC (a PAP2 enzyme) from a *Bacillus subtilis* mutant lacking BacA induces cells to grow with gross morphological distortions and, in some cases, lyse (Zhao *et al.*, 2016). Simultaneous inactivation of LpxE and HupA (PAP2 enzymes) in *Helicobacter pylori* is also lethal (Gasiorowski *et al.*, 2019). Restrictions in Und-PP synthesis or recycling have the same net effect as directly inhibiting Und-PP dephosphorylation. Mutants harboring temperature sensitive alleles of the Und-PP synthase UppS grow misshapen and lyse (Kato *et al.*, 1999, MacCain *et al.*, 2018), as do mutants that trap Und-P in dead-end intermediates (Bhavsar *et al.*, 2001, Cuthbertson *et al.*, 2005, Tatar *et al.*, 2007, Jorgenson *et al.*, 2016, Jorgenson & Young, 2016, Elhenawy *et al.*, 2016). Finally, treating cells (particularly Gram-positive bacteria) with bacitracin, which interferes with Und-PP dephosphorylation through competitive inhibition, is lethal (Smith & Weinberg, 1962).

While *E*. *coli* cells can normally tolerate reductions in Und-PP phosphatase activity (El Ghachi *et al.*, 2004), low Und-P levels provoke strong negative interactions in combination with PG synthase inhibitors. For example, *E*. *coli* cells induced to grow with low Und-P readily lyse in media containing aztreonam or mecillinam, β-lactam antibiotics that inhibit the PG synthesizing activity of penicillin binding proteins (PBPs) (Jorgenson *et al.*, 2019). Similarly, deleting *pgpB* sensitizes cells to cefsulodin, another β-lactam antibiotic (Hernandez-Rocamora *et al.*, 2018). β-lactam antibiotics also synergize with the UppS inhibitor (and fertility drug) clomiphene to kill methicillin-resistant *Staphylococcus aureus* (Farha *et al.*, 2015). Finally, low Und-P levels sensitize *B*. *subtilis* cells to cell wall-active compounds (Lee & Helmann, 2013, Czarny *et al.*, 2014, Peters *et al.*, 2016). Apart from these findings, though, little is known about the connection between Und-P and other pathways. This is due, in part, because most bacteria harbor multiple Und-PP phosphatases. Thus, large-scale genetic interaction studies have likely missed relationships between Und-P metabolism and other pathways due to functional overlap. We therefore conducted a genetic screen to reproduce known Und-P synthetic interactions and to identify new interactions in an *E*. *coli* mutant lacking multiple Und-PP phosphatases. As expected, the screen identified pathways that directly affect Und-PP dephosphorylation but also pathways with no prior connection to Und-P metabolism. These included cell division, DNA replication, signal transduction, and glutathione metabolism. Importantly, some of these connections may prove useful starting points toward developing new combination therapies. An unexpected finding from the screen was the identification of *envZ*, *gor*, *ompR*, and *qseC* as novel morphogenes (i.e., genes required to maintain normal cell shape). Further characterization of *qseC* suggests that phenotypes related to this gene likely result from PG defects.

## Results

### The Δ3PP screen

To identify Und-P synthetic interactions, we designed a screen based on pTB25 (Figure 1A), a derivative of the unstable mini-F plasmid that carries β-galactosidase (*lacZ*) (Bernhardt & de Boer, 2004). In strains that lack the *lac* operon, LacZ produced from pTB25 hydrolyzes X-Gal to form a blue pigment that results in blue colonies on agar plates. Conversely, plasmid-free cells cannot degrade X-Gal and produce white colonies. The plasmid pMAJ95 [P_lac_::*bacA lacZYA*] is a derivative of pTB25 that expresses *lacZ* and the Und-PP phosphatase *bacA* under the control of the *lac* promoter (P_lac_). In MAJ815, an *E*. *coli* strain lacking three Und-PP phosphatases (LpxT, YbjG, and BacA; Δ3PP) and the *lac* operon, pMAJ95 is readily lost as evidenced by the formation of equally sized solid-white and sectored-blue colonies on LB agar containing IPTG and X-Gal (LB-IX) (Figure 1B). Δ3PP cells rely on PgpB for Und-PP phosphatase activity, and a mutant combination lacking all four Und-PP phosphatases (Δ4PP) is not viable (El Ghachi *et al.*, 2005). Indeed, we could only construct Δ4PP cells in the presence of pMAJ95, whose colonies grew solid-blue when plated on LB-IX agar (Figure 1B). This confirms that *bacA* is required for growth in this background.

**Figure 1.**
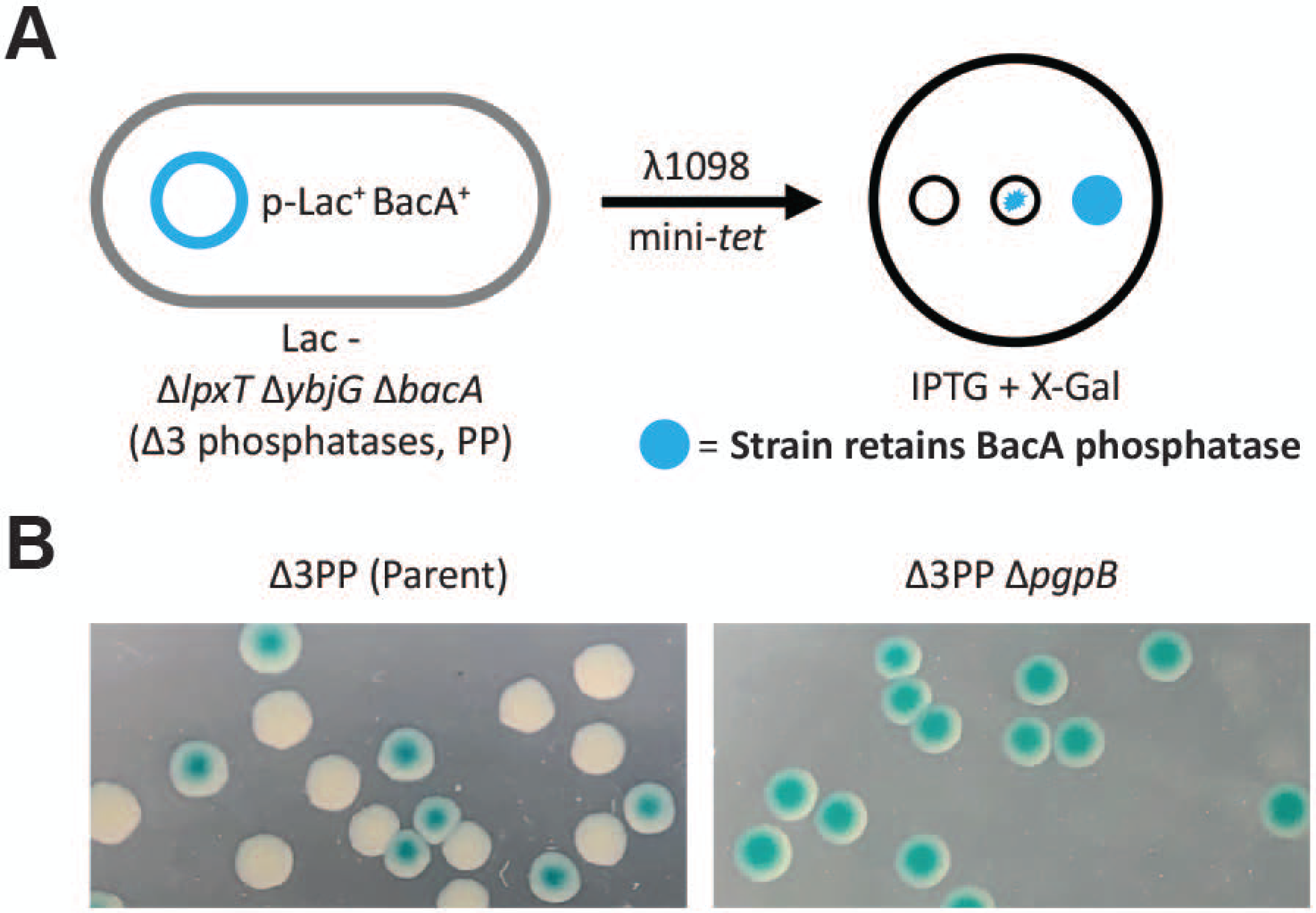
Strategy to screen for Δ3PP synthetic interactions. (A) Δ3PP screen workflow. *E*. *coli* cells lacking three Und-PP phosphatases (Δ*lpxT* Δ*ybjG* Δ*bacA*) and the *lac* operon (Lac -) harbor pMAJ95, a derivative of the unstable mini-F plasmid expressing the *lac* operon and *bacA* in the presence of IPTG. pMAJ95 is readily lost in this background and cells form white or sectored-blue colonies on media containing X-gal. Conversely, introducing synthetically sick or lethal mutant combinations (i.e., λ1098) leads to the retention of pMAJ95 and formation of blue colonies on media containing X-gal. (B) Images depicting white colonies and sectored-blue colonies (left panel) or solid-blue colonies (right panel). The strains tested were MAJ876 (Δ3PP [parent]) and MAJ974 (Δ3PP Δ*pgpB*).

We next mutagenized MAJ815/pMAJ95 by infecting with λ1098 (Way *et al.*, 1984), a λ phage that contains a tetracycline-resistance derivative of Tn*10* (mini-Tn*10*), using a standard λ hop procedure (Way *et al.*, 1984). The mutagenesis yielded approximately 5 × 10^4^ tetracycline resistant mutants that were screened for solid-blue colony formation on LB-IX agar. Since the demand for Und-P increases at higher growth rates (MacCain *et al.*, 2018), this suggested that we might obtain different mutations by plating at different temperatures. Thus, we screened the mutant library at 30°C (∼151,000 colonies), 37°C (∼70,000 colonies), and 42°C (∼83,000 colonies) for solid-blue colony formation. From these, we identified 37 blue colonies that were purified on LB-IX agar. We tested every mutant for solid-blue colony formation at 30°C, 37°C, and 42°C. Upon further characterization, only one of these mutants gave rise to solid-blue colonies on LB-IX agar at every temperature, whose insertion was later mapped to *pgpB* (Table 1). Unexpectedly, for the remaining mutants, we observed the formation of unequally sized solid-blue and solid-white colonies, which we later confirmed (see below). In each case, the blue colonies were significantly larger than the white colonies, and the difference in size between the two became more apparent at higher temperatures. These results demonstrate that ongoing Und-PP phosphatase activity is an important fitness factor.

**Table 1.** *E. coli* genes identified as synthetically sick or lethal in combination with Δ*lpxT* Δ*bacA* Δ*ybjG*.

| Gene name or region | No. of inserts | Location | Product |
| --- | --- | --- | --- |
| Cell wall biogenesis |  |  |  |
| *dacA* | 1 | IM | PG, D,D-carboxypeptidase |
| *ldcA* | 1 | C | PG L,D-carboxypeptidase |
| *mltG* | 1 | OM | PG lytic transglycosylase |
| *P<sub>mreB</sub>* | 1 | IM | Cytoskeletal protein |
| *mrcB* * | 1 | IM | PG bifunctional synthase |
| *pgpB* * | 1 | IM | Und-PP phosphatase |
| *pal* | 2 | OM | PG associated OM lipoprotein |
| *tolA* | 1 | IM | Cell envelope integrity IM protein |
| *tolB* | 1 | IM | Cell envelope integrity IM protein |
| Cell division |  |  |  |
| *ftsK* | 1 | IM | Cell division DNA translocase |
| *minD* | 1 | C | FtsZ ring positioning protein |
| *T<sub>minE</sub>* | 1 | C | FtsZ ring positioning protein |
| Disulfide bond formation |  |  |  |
| *gor*, *P<sub>gor</sub>* | 2 | C | Glutathione reductase |
| ECA synthesis |  |  |  |
| *wecD* | 1 | C | dTDP-fucosamine acetyltransferase |
| *wecE* | 1 | C | dTDP-4-dehydro-6-deoxy-D-glucose transaminase |
| *wzxE* | 4 | IM | ECA lipid III flippase |
| Replication and Repair |  |  |  |
| *rep* | 1 | C | DNA helicase |
| *seqA* | 1 | C | Negative regulator DNA replication |
| *topA* | 1 | C | DNA topoisomerase 1 |

| Signal transduction |  |  |  |
| --- | --- | --- | --- |
| *cpxR* | 1 | C | Response regulator for CpxA |
| *envZ* | 1 | IM | Sensor protein for OmpR |
| *ompR* | 1 | C | Response regulator for EnvZ |
| *qseC* | 2 | IM | Sensor protein for QseB |
| Other |  |  |  |
| *lon* | 1 | C | Protease |

Abbreviations: C, cytosol; IM, inner membrane; OM, outer membrane; ECA, enterobacterial common antigen; P, promoter; T, terminator.
\*Synthetic lethal. All other gene combinations are synthetic sick.

Mapping mini-Tn*10* insertions by arbitrary polymerase chain reaction (PCR) (O’Toole & Kolter, 1998) uncovered 30 unique insertions in 22 genes and 3 intergenic regions (Table 1). Categorizing these genes and loci (Table 1) by function revealed that the majority of insertions (33%) were in genes that directly affect cell wall metabolism. These included insertions in cell wall hydrolases (*dacA*, *ldcA*, and *mltG*), cell wall synthases (*mrcB*), the Tol-Pal system (*tolA*, *tolB*, and *pal*), and the Und-PP encoding phosphatase *pgpB*, whose identification served as a positive proof-of-principle for the Δ3PP screen. We also isolated an insertion in the promoter of the actin homolog *mreB* (P*_mreB_*) (Table S1). Genes required for the late stages of enterobacterial common antigen (ECA) synthesis were also identified (*wecD*, *wecE*, and *wzxE*). Since disruptions in the ECA pathway sequester Und-PP-linked intermediates (Jorgenson *et al.*, 2016), thus preventing Und-P(P) recycling, we expected to uncover this class of mutants. The remaining insertions were in genes or regions with no obvious connection to Und-P metabolism. These included genes required for cell division (*ftsK*), glutathione formation (*gor*), DNA replication and repair (*rep*, *seqA*, and *topA*), and signal transduction (*cpxR*, *envZ*/*ompR*, and *qseC*). We note that insertions in *ftsK* and *topA* were outside the essential region of these genes (Table S1) (Goodall *et al.*, 2018).

To verify that the colony phenotypes were due to the inactivation of these genes alone, we generated null mutations for most non-essential genes identified in the Δ3PP screen in the MAJ815/pMAJ95 background. A representative mutant was picked for the ECA biosynthesis (Δ*wzxE*) and Tol-Pal (Δ*pal*) pathways. In every case, the null mutants mirrored the colony phenotypes of their corresponding transposon mutant. Therefore, the null mutants were used in all subsequent studies. We also reintroduced the mini-Tn*10* insertion in P*_mreB_* into MAJ815/pMAJ95 by P1 transduction and confirmed the colony phenotype (Figure 2B). As stated previously, only loss of PgpB resulted in solid-blue colonies at all temperatures (Figure 2), an indication of synthetic lethality. Alternatively, we observed that PBP1B (Δ*mrcB*) and P*_mreB_* mutants gave rise to exclusively solid-blue colonies at 42°C (Figure 2B). The outer membrane lipoprotein LpoB is required to activate PBP1B (Typas *et al.*, 2010, Paradis-Bleau *et al.*, 2010), but was not identified in the Δ3PP screen. Therefore, we determined whether a mutant lacking LpoB mirrors the phenotype a PBP1B-null mutant in the Δ3PP background. As expected, Δ3PP Δ*lpoB* cells produced solid-blue colonies only at 42°C (Figure S1). These results demonstrate that the screen was not saturating at 42°C. Removing *ompR* or *wzxE*, as well as disrupting *topA* in Δ3PP cells was also deleterious at high temperature, resulting in vanishingly few white colonies at 42°C (Figure 2C). Since mutants lacking Lon or Min filament/produce minicells, we confirmed their transposon mutants by microscopy (see below). In total, the Δ3PP screen identified system wide connections to Und-P metabolism.

**Figure 2.**
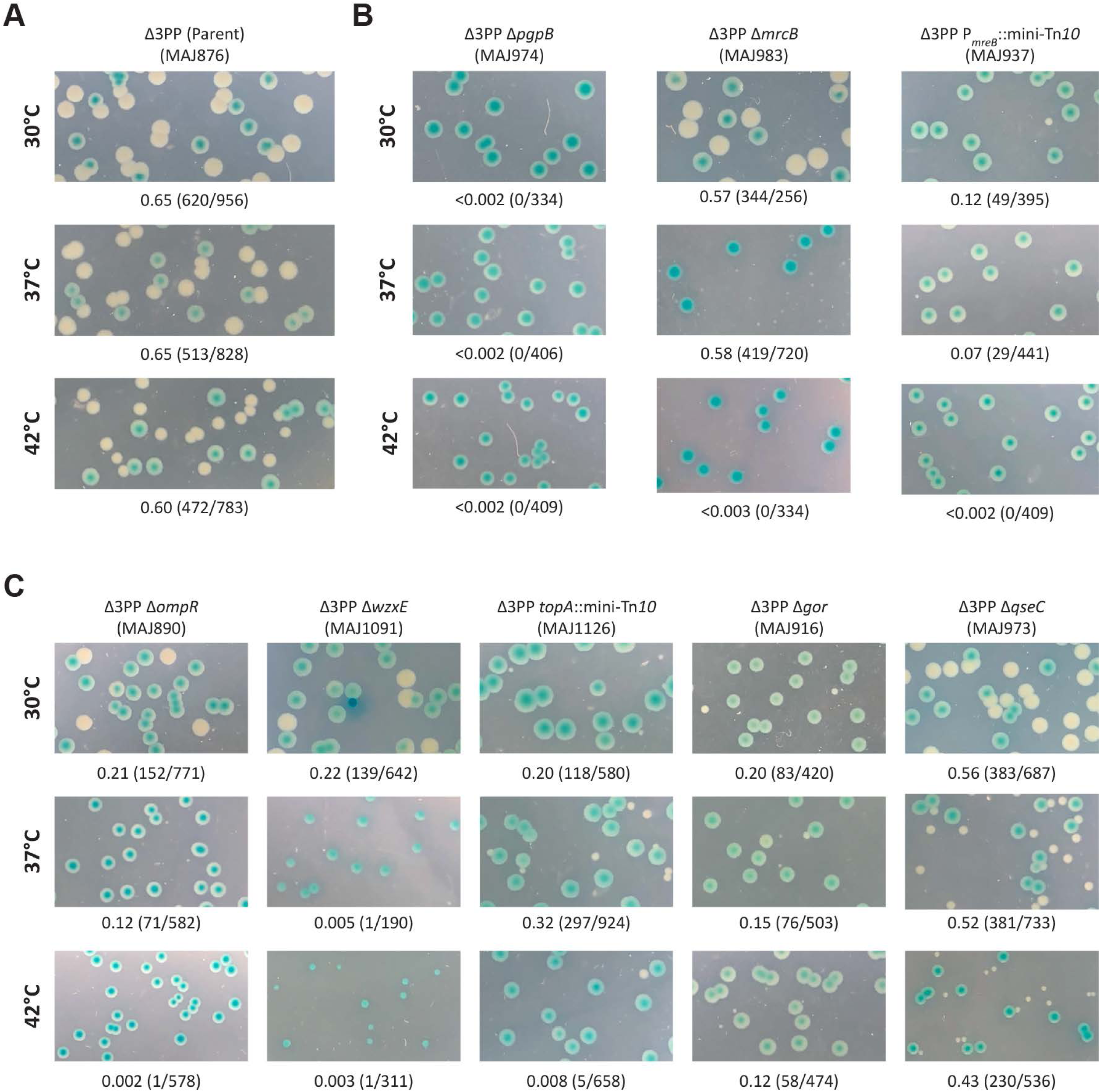
Viability of Δ3PP mutant derivatives. (A) pMAJ95 readily segregates in the Δ3PP parent, resulting in white or sectored-blue colonies at 30°C, 37°C, and 42°C. (B and C) pMAJ95 is retained in mutants that are synthetically lethal in the Δ3PP background and colonies appear solid-blue. Alternatively, the frequency of pMAJ95 segregation is reduced in mutants that are synthetically sick in the Δ3PP background because pMAJ95 confers a growth advantage. Above photographs: genotype and strain designation. Below photographs: fraction of white colonies to the total number of colonies. Additional Δ3PP mutant derivatives are shown in Figure S1 in the supplemental material.

### The Δ3PP screen uncovers known and novel morphological determinants

A steady supply of Und-P is required to maintain normal cell shape (Jorgenson *et al.*, 2016, Jorgenson & Young, 2016), which suggested that synthetic mutations in the Δ3PP background might provoke shape defects. To determine whether our Δ3PP mutant derivatives harbored cell shape defects, cells were cultured in LB medium with or without chloramphenicol (to select for pMAJ95) and IPTG (to express *bacA*) at 37°C and 42°C. While loss of *bacA* expression had little effect on the morphology for the Δ3PP parent at 37°C (Figure S3A) or 42°C (Figure 3A), *bacA* expression was required to maintain cell shape and integrity for all Δ3PP mutants (Figure 3, S2, and S3), especially at 42°C (Figure 3 and S2B; compare the morphologies for +p*bacA* to −p*baA*). We broadly categorized the phenotypes we observed as cell division or morphogenesis-related. All mutants were imaged at 37°C (Figure S2 and S3) and 42°C (Figure 3 and S2).

**Figure 3.**
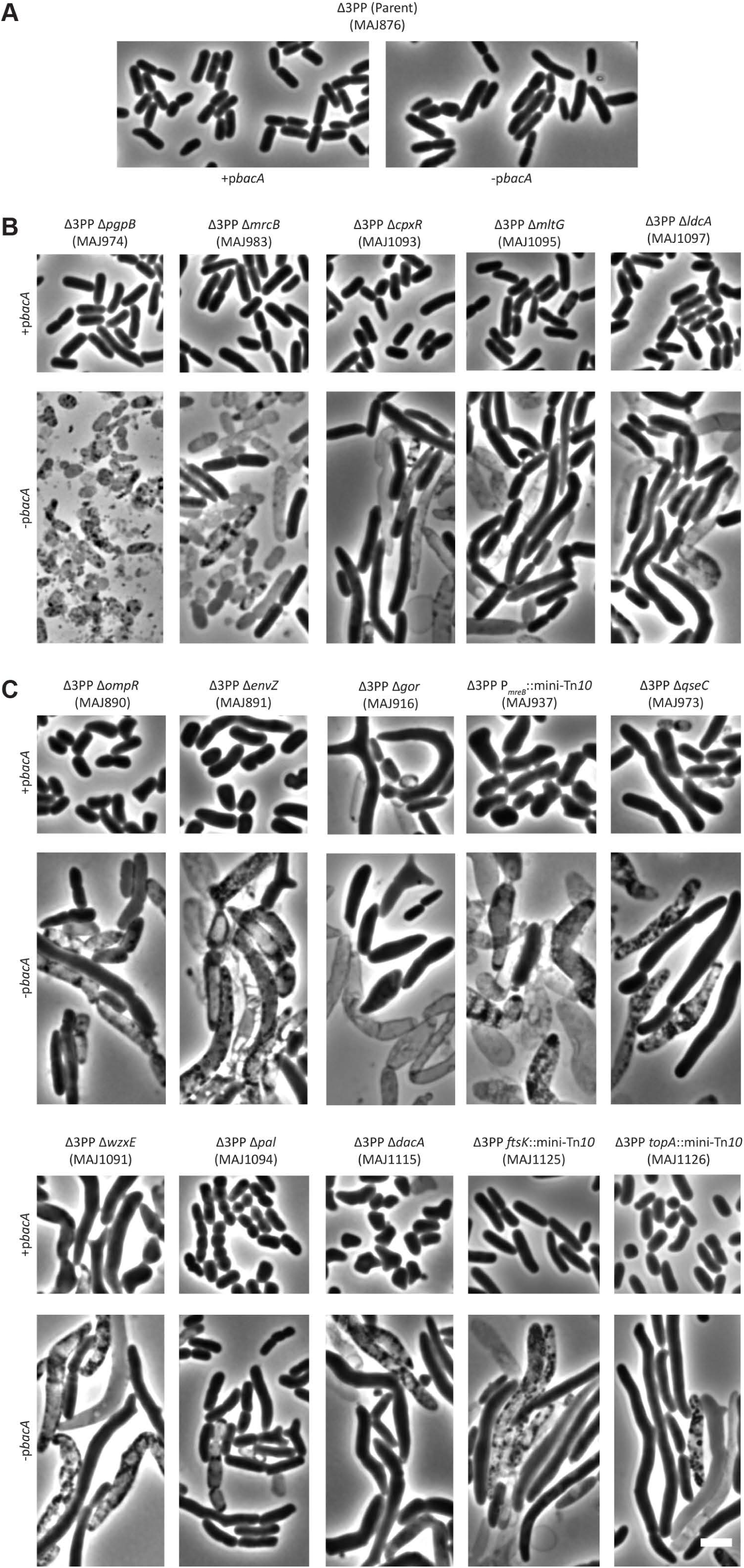
Morphology of Δ3PP mutant derivatives at 42°C. Cells with the indicated genotypes were grown at 42°C in LB (−p*bacA*) or LB containing chloramphenicol and 500 µM IPTG (+p*bacA*) until the culture reached an OD_600_ of 0.3-0.4. The cells were then photographed by phase-contrast microscopy. Images of Δ3PP mutants grown at 37°C are shown in Figure S3 in the supplemental material. The white bar represents 3 µm. (A) The Δ3PP parent. (B) Mutant derivatives that produce shape defects in the absence of *bacA* expression. (C) Mutant derivatives that produce shape defects independent of *bacA* expression. The data is representative of at least two independent experiments.

Mutants affected for cell division grew as long filaments irrespective of *bacA* expression (Figure S2). This included the Lon protease, which is required to degrade SulA, an inhibitor of FtsZ (which initiates cell division) ring formation (Mizusawa & Gottesman, 1983, Higashitani *et al.*, 1997) (Figure S2). Since SulA filaments cells in the absence of Lon (Adler & Hardigree, 1964, Gayda *et al.*, 1976), we reasoned that SulA was also responsible for filamenting Δ3PP *lon*::mini-Tn*10* cells (Figure S2). We note, however, that SulA is thermolabile and aggregates at 42°C (Mizrahi *et al.*, 2007). Thus, it is unlikely that SulA is soley responsible for the shape defect of Δ3PP *lon*::mini-Tn*10* cells at 42°C. Mutations that affect *seqA* (Lu *et al.*, 1994, Pedersen *et al.*, 2017) or *rep* (Michel *et al.*, 1997) lead to double-stranded DNA breaks, which activates the SOS response (Ossanna & Mount, 1989, Rotman *et al.*, 2014), triggering *sulA* expression and thus filamentation (Figure S2). The Min proteins MinC, MinD, and MinE function to restrict FtsZ ring formation to midcell (de Boer *et al.*, 1989). Importantly, loss of MinE enables MinCD complexes to inhibit FtsZ-ring formation throughout the cell, causing filamentation (de Boer *et al.*, 1989). This data suggests that T*_minE_*::mini-Tn*10* cells filament due to limiting amounts of MinE (Figure S2). We also note that mutants that lack the DNA translocase FtsK grow as long filaments (Begg *et al.*, 1995). However, an insertion outside the essential region of FtsK (Table S1) (Goodall *et al.*, 2018) only had a minor effect on cell length in Δ3PP cells when BacA was present (Figure 3C, +p*bacA*). Interestingly, all of the filamenting mutants lysed in the absence of *bacA* expression when grown at 42°C (Figure S2B). This suggests that ongoing Und-PP phosphatase activity is critical in fast growing cells that are blocked for cell division. More generally, we interpret the isolation of mutants that filament to issues pertaining to plasmid segregation. Since replication of the unstable plasmid and the chromosome initiate simultaneously (Frame & Bishop, 1971), this means that most, if not all, cell filaments will contain multiple copies of the mini-F plasmid due to the presence of multiple chromosomes. Thus, mutants that filament are more likely to inherit the unstable plasmid at cell division.

For those mutants affected for morphogenesis, we further divided these phenotypes as BacA-dependent or BacA-independent. BacA-dependent mutants produced rod-shaped cells when expressing *bacA,* but grew misshapen and/or lysed in its absence (Figure 3B; see also Figure S3B for 37°C morphologies). This mutant class was composed entirely of genes that affect PG metabolism, either directly or indirectly. Factors that directly affect PG metabolism included the Und-PP phosphatase PgpB (El Ghachi *et al.*, 2005), the cell wall synthase PB1B (Δ*mrcB*) (Sauvage *et al.*, 2008), and the cell wall degrading enzymes MltG (Yunck *et al.*, 2016) and LdcA (Templin *et al.*, 1999). As expected, cells lacking all four Und-PP phosphatases (Δ*pgpB*) lysed completely, as did a significant fraction of Δ3PP cells deleted for *mrcB* (Figure 3B). This confirms and extends our previous finding that simultaneously inhibiting the production of Und-P and PG metabolism produces a synergistic response leading to cell lysis (Jorgenson *et al.*, 2019). We also identified the response regulator CpxR from the CpxAR two-component signal transduction system (Raivio, 2014), which indirectly affects PG metabolism by regulating the expression of PG modifying enzymes (Weatherspoon-Griffin *et al.*, 2011, Bernal-Cabas *et al.*, 2015). This suggests that the Cpx response is critical for viability when Und-P is limiting.

BacA-independent mutants produced morphological defects, even when expressing *bacA* (Figure 3C; see also Figure S3C for 37°C morphologies). Again, many of the genes disrupted in this class of mutants produce factors already known to maintain cell shape. This included the ECA flippase WzxE (Rick *et al.*, 2003) (Figure 3C), whose absence sequesters Und-P in non-recyclable intermediates causing cells to grow misshapen (Jorgenson *et al.*, 2016). In addition, mutants lacking Pal do not efficiently constrict the cell envelope and produce chains of unseparated cells (Gerding *et al.*, 2007), which we observed at 42°C (Figure 3C). DacA cleaves the terminal D-alanine from PG peptide side chain (Spratt & Strominger, 1976, Matsuhashi *et al.*, 1979) and mutants grow as branched cells (Nelson & Young, 2000), which we also observed (Figure 3C and S3C). The *mre* locus encodes MreBCD and are required to maintain rod shape (Wachi *et al.*, 1989, Wachi *et al.*, 1987) and MreB requires a steady supply of Und-P for filament formation (Schirner *et al.*, 2015). Not surprisingly, cells with an insertion in the promoter region of *mreB* were misshapen (Figure 3C and S3, +p*bacA*) and lysed in the absence of BacA (Figure 3C and S3, −p*bacA*). Finally, TopA relaxes supercoiled DNA (Kirkegaard & Wang, 1985) and null mutants (harboring compensatory mutations) form long filaments (Usongo & Drolet, 2014); however, an insertion in the nonessential region of *topA* caused cells to grow as misshapen rods and spheres (Figure 3C and S3C).

The BacA-independent mutants also contained several genes whose products were not known to affect morphology. This included both components of the EnvZ/OmpR two-component system, the glutathione reductase Gor, and QseC, the sensor kinase component of the QseBC two-component system. To determine whether these shape defects were due to the loss of these genes alone (and not due to the combined loss of LpxT and YbjG), we deleted *envZ*, *gor*, *ompR*, or *qseC* from the parental wild type (i.e., MG1655 Δ*lacIZYA*). Importantly, these mutants exhibited morphologies similar to their Δ3PP derivatives when evaluated by microscopy and flow cytometry (compare morphologies in Figure 4 to those in Figure 3C and S3C). A closer inspection revealed that cells lacking QseC were 30% longer and 15% wider than the parental wild type when cultured at 37°C and 54% longer and 27% wider at 42°C (Table 2). When examined by flow cytometry, the distribution of the forward scatter area of Δ*qseC* cells was shifted to the right when compared to that of the parental wild type (Figure 4B), confirming that Δ*qseC* cells are enlarged. Mutants lacking EnvZ, Gor, or OmpR were also enlarged (Figure 4), but we also observed branching in these mutant backgrounds (Figure 4A). As expected, expressing these genes from a plasmid reversed these shape defects (Figure S4). In summary, disrupting Und-P metabolism triggers shape defects in mutants whose gene products have direct and tangential connections to morphogenesis.

**Figure 4.**
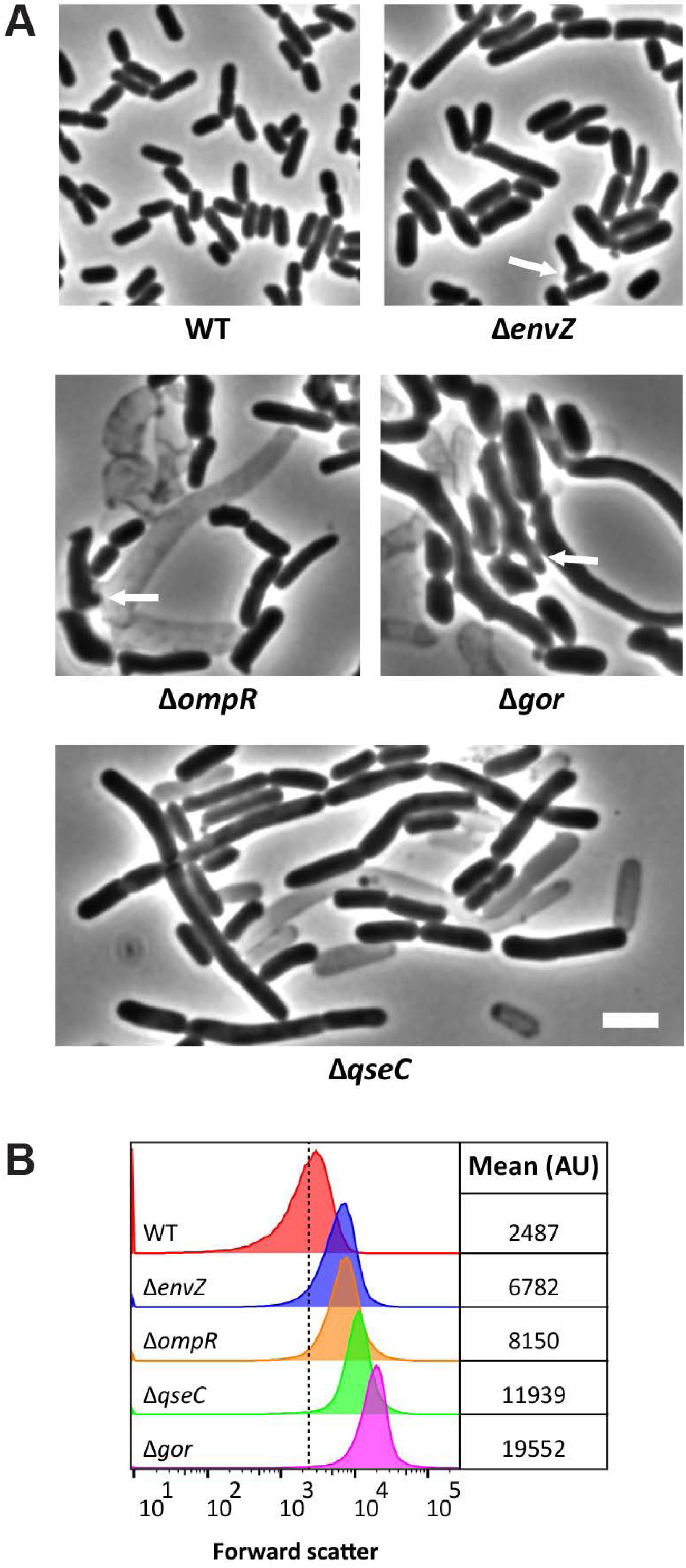
New shape-determinants. (A) Cells with the indicated genotypes were grown at 37°C in LB until the culture reached an OD_600_ of 0.5. The cells were then photographed by phase-contrast microscopy. White arrows point to examples of branching. The white bar represents 3 µm. (B) Live cells from panel A were also examined by flow cytometry. Histograms of the forward scatter area from 100,000 events (cells) are shown. The mean cell size of the wild type is represented by the dashed line and is expressed in arbitrary units (AU). Data are representative of two independent experiments. The strains shown are MAJ25 (WT), MAJ964 (Δ*envZ*), MAJ938 (Δ*ompR*), MAJ969 (Δ*gor*), and MAJ1005 (Δ*qseC*).

**Table 2.** Morphological phenotypes of an *E. coli qseC* null mutant.

| Genotype<sup>a</sup> | Temp (°C) | No. of cells evaluated | Avg. length, μm, (SD) | Avg. width, μm (SD) | % cells with indicated no. of constrictions | | |
| --- | --- | --- | --- | --- | --- | --- | --- |
|  |  |  |  |  | 0 | 1 | >1 |
| WT | 30 | 371 | 2.9 (0.7) | 1.2 (0.1) | 68 | 32 | 0 |
|  | 37 | 319 | 2.6 (0.6) | 1.2 (0.1) | 62 | 38 | 0 |
|  | 42 | 261 | 2.6 (0.7) | 1.3 (0.2) | 60 | 40 | 0 |
| $\Delta qseC$ | 30 | 196 | 3.4 (1.1) | 1.3 (0.2) | 73 | 27 | 0 |
|  | 37 | 341 | 3.5 (1.2) | 1.4 (0.2) | 62 | 38 | 0 |
|  | 42 | 264 | 4.5 (2.7) | 1.7 (0.2) | 58 | 42 | 0 |

<sup>a</sup>Strains used were MAJ25 (WT) and MAJ1005 ( $\Delta qseC$ ).

### QseB overactivation disrupts cell shape

At this point, we decided to characterize the Δ*qseC* mutant and will report on the *envZ*, *gor*, and *ompR* mutants later. QseC is a sensor kinase that phosphorylates QseB in response to autoinducer-3, epinephrine, and norepinephrine (Clarke *et al.*, 2006). Phosphorylated QseB positively regulates its expression (Clarke & Sperandio, 2005a) as well the expression of *flhDC*, flagellar synthesis genes (Clarke & Sperandio, 2005b). Loss of QseC causes cross-phosphorylation of its response regulator QseB through PmrB (Guckes *et al.*, 2013), another sensor kinase, resulting in constitutive action of QseB (Figure 5A). Overactivation of QseB disrupts metabolism (Hadjifrangiskou *et al.*, 2011), reduces expression of virulence-associated genes (Kostakioti *et al.*, 2009), and (now) disrupts cell shape (Figure 4). Importantly, Δ*qseC* phenotypes are suppressed by deleting *qseB* or *pmrB* (Kostakioti *et al.*, 2009, Bearson *et al.*, 2010, Guckes *et al.*, 2013). To determine if the shape defect of Δ*qseC* cells was related to QseB overactivation, we first constructed a mutant lacking QseC and QseB. Strikingly, loss of QseB completely reversed the shape defect of Δ*qseC* cells when grown in LB medium at 42°C (Figure 5B and 5C). Next, we determined whether deleting *pmrB* in Δ*qseC* cells would have a similar effect. Interestingly, loss of PmrB only partially reversed the shape defect of Δ*qseC* cells (Figure 5B and 5C). This result suggests that additional factors are capable of activating QseB. The PmrAB two-component system also positively regulates *qseB* expression at the level of transcription through the PmrA response regulator, and deleting *pmrA* from Δ*qseC* cells can suppress some but not all Δ*qseC* phenotypes (Guckes *et al.*, 2013). In this case, deleting *pmrA* had no effect on the shape of Δ*qseC* cells (Figure S5). Collectively, these findings demonstrate that constitutive activation of QseB, likely by multiple kinases, results in cell shape defects.

**Figure 5.**
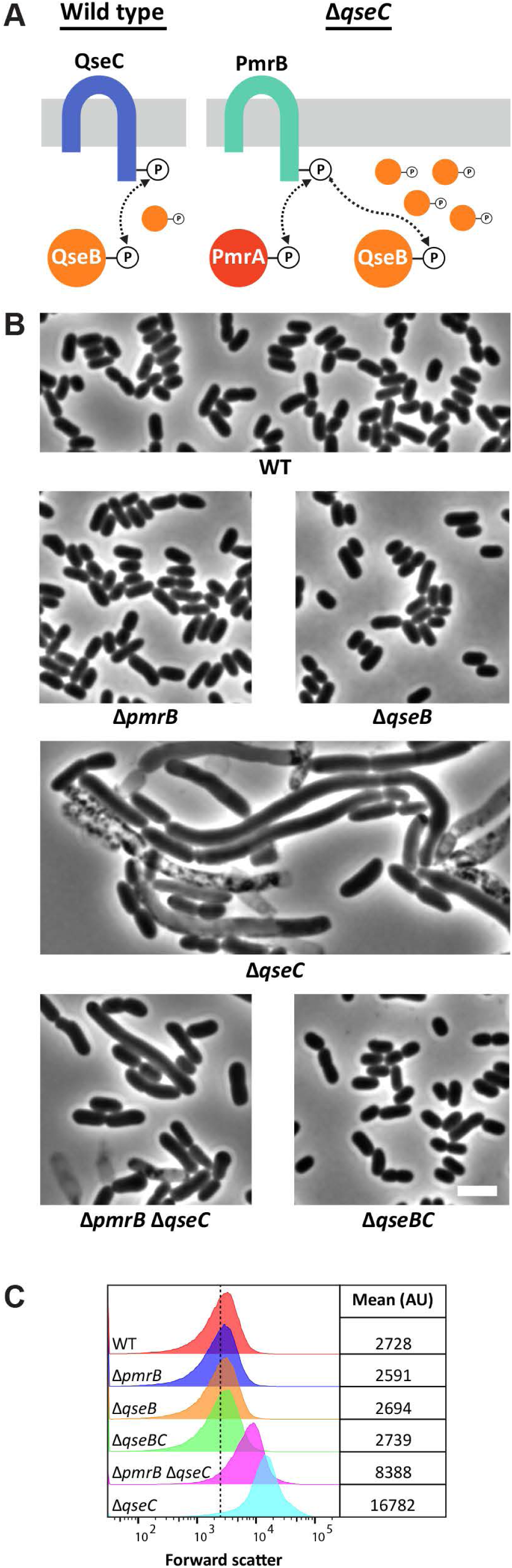
QseB overactivation disrupts cell shape. (A) Schematic depicting activated QseC phosphorylating and activating QseB (<u>wild type</u>). PmrB phosphorylates QseB in the absence of QseC (<u>Δ*qseC*</u>). Unlike QseC, however, PmrB cannot efficiently dephosphorylate QseB, leading to excessive amounts of activated QseB (Guckes *et al.*, 2013). (B) Cells with the indicated genotypes were grown at 42°C in LB until the culture reached an OD_600_ of 0.5. The cells were then photographed by phase-contrast microscopy. The white bar represents 3 µm. (C) Live cells from panel B were also examined by flow cytometry. Histograms of the forward scatter area from 100,000 events (cells) are shown. The mean cell size of the wild type is represented by the dashed line and is expressed in arbitrary units (AU). Data are representative of two independent experiments. The strains shown are MAJ25 (WT), MAJ1002 (Δ*pmrB*), MAJ962 (Δ*qseB*), MAJ1005 (Δ*qseC*), MAJ1003 (Δ*pmrB* Δ*qseC*), and MAJ999 (Δ*qseBC*).

### Evidence ruling out YgiW in the shape defect of Δ*qseC* cells

In *E*. *coli* UTI189, QseB overactivation was observed to alter the expression of 443 genes (Hadjifrangiskou *et al.*, 2011). Of the genes that were highly altered (e.g., *qseB*), the most upregulated gene was *ygiW* (fold change 2.8 × 10^17^). Interestingly, *ygiW* is located directly upstream of *qseBC* (Hadjifrangiskou *et al.*, 2011) and codes for a predicted periplasmic protein of the bacterial oligonucleotide/oligosaccharide binding-fold (BOF, PF04076) protein family (Ginalski *et al.*, 2004). Notably, the YgiW homolog in *Salmonella typhimurium* (i.e., VisP) binds PG in a pull-down assay (Moreira *et al.*, 2013). Based on these observations, we reasoned that *ygiW* overexpression may interfere with PG synthesis in Δ*qseC* cells. However, deleting *ygiW* did not reverse the shape defect of Δ*qseC* cells (Figure S6). This result indicates that *ygiW* has no effect on shaping Δ*qseC* cells.

### Multicopy suppression of the shape defect produced by Δ*qseC* cells

To further determine the morphological basis for the Δ*qseC* shape defect, we isolated multicopy plasmids that reverse the shape defect of cells lacking QseC. Since Δ3PPΔ*qseC* cells lacking pMAJ95 (i.e., *bacA*) produce small-colonies (Figure 2C) and grow as misshapen cells (Figure 3C and S3C), we reasoned that multicopy suppressing plasmids would produce large-colonies that grow as rod-shaped cells (Figure 6A). To do this, we first introduced a pBR322-based *E*. *coli* library (Ulbrandt *et al.*, 1997) into strain MAJ973 (Δ3PPΔ*qseC*/pMAJ95), and screened approximately 30,000 colonies on LB-IX ampicillin (ampicillin selects for pBR322 derivatives) plates at 42°C for light blue colony formation, indicating segregation of pMAJ95. From these, we identified 21 light blue colonies that produced large colonies when purified on LB-IX agar at 42°C and rod-shaped cells when grown in LB medium containing ampicillin and 500 µM IPTG at 42°C. Conversely, small-colony white variants always produced misshapen cells. Candidate plasmids were then isolated and transformed into Δ*qseC* (MAJ1005) cells and examined for shape suppression at 42°C as above. From these, we identified two classes of suppressing plasmids. The first class fully restored Δ*qseC* cells to the size and shape of wild type cells, and included plasmids harboring *qseC* or *rprA*, a small regulatory RNA (Majdalani *et al.*, 2001). The second class of suppressing plasmids partially reversed the shape defect of Δ*qseC* cells, and consisted entirely of plasmids containing *pdhR*, a transcriptional regulator (Quail & Guest, 1995). Plasmids expressing only *qseC*, *rprA*, or *pdhR* produced the same results as the pBR322 derivatives (Figure 6 and S7), confirming that overexpressing *rprA* and, to a lesser extent, *pdhR* can suppress the shape defect of Δ*qseC* cells. To our knowledge, multicopy shape suppression by *rprA* is the first example of a small RNA acting as a shape suppressor. During the course of these experiments, we also tested whether overexpressing the Und-PP synthase *uppS* could suppress the shape defect of Δ*qseC* cells. However, *uppS* overexpression had no effect on the shape of Δ*qseC* cells (Figure S4). This suggests Und-P(P) is not limiting in QseC cells.

**Figure 6.**
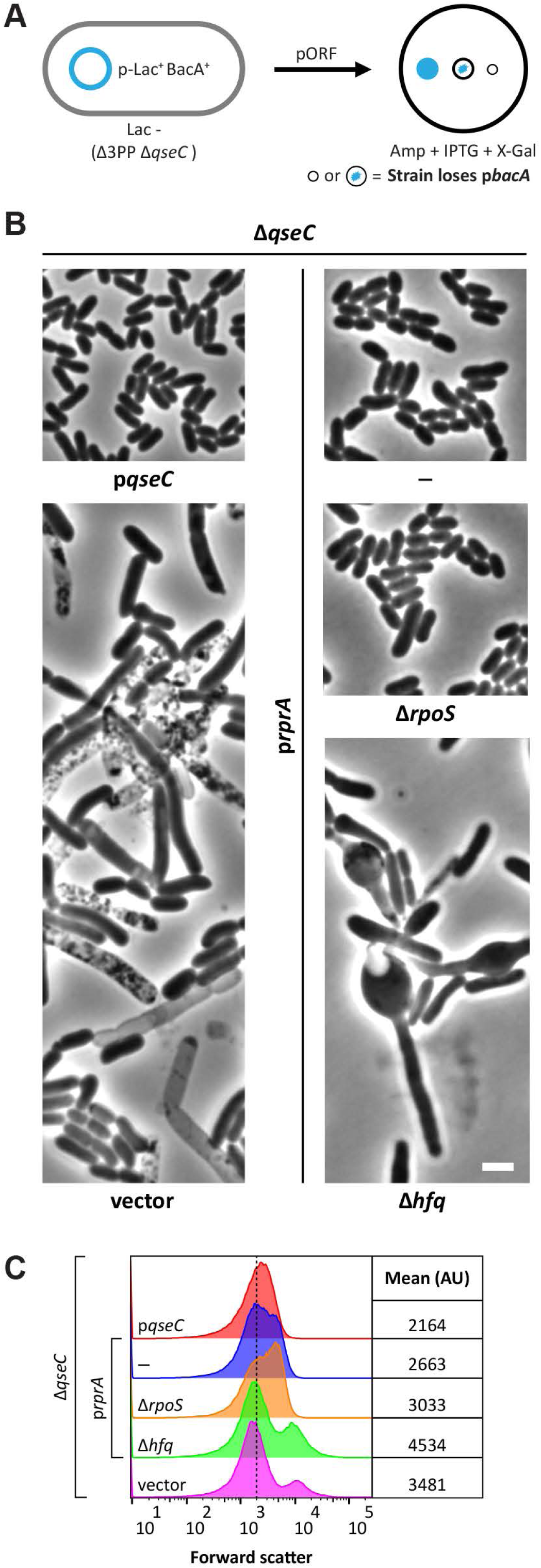
Suppression of Δ*qseC* shape defects. (A) Workflow of strategy to uncover multicopy plasmids that suppress the shape defect of Δ*qseC* cells. Δ3PP Δ*qseC* cells harboring pMAJ95 produce predominantly blue colonies at 42°C (cf., Figure 2C). However, pMAJ95 segregates in cells containing plasmids (pORF) that suppress the Δ*qseC* shape defect, and appear light blue (i.e., sectored-blue). Note that white colonies (indicating loss of pMAJ95) always yielded misshapen cells. (B) Micrographs of Δ*qseC* cells containing derivatives of pBRplac that express *qseC* or the small RNA *rprA*. Cells with the indicated genotypes were grown at 42°C in LB until the culture reached an OD_600_ of 0.5. The cells were then photographed by phase-contrast microscopy. The white bar represents 3 µm. (C) Live cells from panel B were also examined by flow cytometry. Histograms of the forward scatter area from 100,000 events (cells) are shown. The mean cell size of the wild type is represented by the dashed line and is expressed in arbitrary units (AU). Data are representative of two independent experiments. The strains shown are MAJ1177 (p*qseC*), MAJ1143 (vector), MAJ1144 (—), MAJ1151 (Δ*rpoS*), and MAJ1163 (Δ*hfq*/p*rprA*).

### Shape suppression of Δ*qseC* cells by multicopy *rprA* requires Hfq but not RpoS

We next sought to determine what factors were required for shape suppression of Δ*qseC* cells by multicopy *rprA*. RprA (RpoS regulator RNA) positively regulates translation of the stationary phase sigma factor RpoS (Majdalani *et al.*, 2001) by binding to and stabilizing *rpoS* mRNA (Majdalani *et al.*, 2002), an interaction that is enhanced by the RNA-binding protein Hfq (Soper *et al.*, 2010). We reasoned that the function of one of these factors was important to suppress the shape defect of Δ*qseC* cells by multicopy *rprA*. To determine the contribution of RpoS to multicopy shape suppression of Δ*qseC* cells by RprA, we overexpressed *rprA* in a Δ*qseC* Δ*rpoS* mutant. Surprisingly, loss of RpoS had no appreciable effect on shape suppression (Figure 6B and 6C). This suggests that activation of the general stress response is not required to suppress the shape defect of Δ*qseC* cells by multicopy *rprA*. Since Hfq enhances RprA-RNA binding (Soper *et al.*, 2010), we reasoned that loss of Hfq might prevent RprA activity. Indeed, multicopy *rprA* failed to restore cell shape to Δ*qseC* Δ*hfq* cells, which formed bulges (Figure 6B and 6C). This result suggests an alternative RprA-RNA interaction is required to suppress the shape defect of Δ*qseC* cells. However, we note that Hfq is required to maintain normal cell shape (Tsui *et al.*, 1994, Lybecker *et al.*, 2010, Bai *et al.*, 2010, Wilms *et al.*, 2012, Boudry *et al.*, 2014, Irnov *et al.*, 2017) (Figure S5). Thus, we cannot conclusively state whether *rprA* suppression requires Hfq (though we consider it likely). RprA also regulates the expression of *csgD* and *ydaM* (Jorgensen *et al.*, 2012, Mika *et al.*, 2012), but these factors were not required for multicopy shape suppression of Δ*qseC* cell by RprA (Figure S8). Collectively, these data indicate that RprA plays a role in maintaining cell shape by an unknown mechanism.

## Discussion

Walled bacteria employ polyprenyl carrier lipids to assemble and transport cell wall intermediates (Manat *et al.*, 2014). In *E*. *coli*, like most bacteria, Und-P serves as the primary lipid carrier and is essential (Kato *et al.*, 1999). Since Und-P originates from the dephosphorylation of Und-PP, the phosphatases required to generate the monophosphate linkage are conditionally essential (El Ghachi *et al.*, 2004, El Ghachi *et al*., 2005). Here, we took advantage of this functional overlap to identify new connections to Und-P metabolism. Utilizing a derivative of the unstable mini-plasmid system (Bernhardt & de Boer, 2004), we identified synthetically sick and lethal interactions in the absence of three (out of four) Und-PP phosphatases in *E*. *coli*. The screen was highly successful, netting several known connections to Und-P metabolism as well as entirely new connections. The absolute requirement of Und-P for PG metabolism (Kato *et al.*, 1999) means that the screen was primed to uncover PG-intersecting pathways, which we show were present in greater numbers than previously thought. In short, we demonstrate the utility of disrupting Und-P metabolism to find new ways to inhibit bacterial growth and uncover subtle biological circuits.

### Und-PP metabolism questions not resolved by the Δ3PP screen

Several open questions regarding Und-P(P) metabolism were not resolved by the Δ3PP screen. For example, Und-PP dephosphorylation occurs extracellularly (Tatar *et al.*, 2007, Touze *et al*., 2008, Manat *et al.*, 2015), yet it is not known how Und-PP is flipped from the inner side of the cytoplasmic membrane to the outer side or how it is flopped back as Und-P. In addition, it is not known whether there are additional Und-PP phosphatases, noting that the BacA and PAP2 protein families share no sequence similarity (El Ghachi *et al.*, 2005). Finally, whether all Und-P-utilizing pathways have been identified in *E*. *coli* is not settled. The results from the Δ3PP screen suggest two possibilities. First, factors that transport, dephosphorylate, or utilize Und-P(P) are already known, to varying degrees. We consider this the most likely possibility. Alternatively, additional factors required for Und-PP metabolism exist but are essential or were missed for other (unknown) reasons. Identifying Und-P(P) interactions in environmental isolates or Gram-positive organisms may help distinguish between these possibilities. More broadly, questions regarding the synthesis and metabolism of Und-P(P) have important implications toward treating bacterial infections (see below) and engineering bacteria to produce isoprenoid derivatives for high-value pharmaceuticals and industrial products (Tetali, 2019).

### Surface area to volume requirements likely explain the basis for Δ3PP mutant morphologies

Why do Δ3PP mutant derivatives grow bigger in the absence of BacA? While it makes some sense that inhibiting multiple PG-affecting pathways would cause cells to grow more misshapen, it is not obvious why they always grow larger. This basic question is rooted in fundamental laws governing how cells grow and determine their shape. The ‘relative rates’ model assumes that bacteria alter their dimensions so that the ratio of surface area to volume (SA/V) homeostasis equals the ratio of (β) surface synthesis to (α) volume synthesis (i.e., SA/V=β/α) (Harris *et al.*, 2014, Harris & Theriot, 2018, Harris & Theriot, 2016). Thus, the ‘relative rates’ model predicts that in cases where surface synthesis (β) decreases and volume synthesis (α) increases, cells will reduce SA/V to re-establish SA/V homeostasis. This type of situation occurs when PG precursor synthesis is inhibited (Harris & Theriot, 2016, Si *et al.*, 2017, Real & Henriques, 2006). In such cases, bacteria reduce SA/V by growing longer and wider (i.e., bigger). Since decreasing Und-PP dephosphorylation is expected to reduce PG precursor formation, the ‘relative rates’ model would predict Δ3PP cells to grow even longer and wider if inhibited further for PG synthesis, which we observed for all Δ3PP mutant derivatives (Figure 3 and S3). In cases where cells lysed, dimensional changes were likely insufficient to lower SA/V beyond the threshold needed to withstand the force generated by internal osmotic (turgor) pressure. In short, the ‘relative rates’ model provides a likely explanation why inhibiting Und-PP dephosphorylation primes cells with otherwise imperceptible defects in PG synthesis grow larger. Whether reducing PG precursors in other contexts would identify similar pathways is unknown.

### Reexamining the physiological functions of the QseBC two-component system

The QseBC system is credited as a global regulator (Weigel & Demuth, 2016), responsible for controlling biofilm formation and virulence. This designation is due, in large measure, to the use of *qseC* mutants. For example, loss of QseC impairs biofilm formation of *Aggregatibacter actinomycetemcomitans* (Novak *et al.*, 2010, Juarez-Rodriguez *et al.*, 2013) and *Haemophilus influenzae* (Unal *et al.*, 2012). Similarly, *qseC* mutants are attenuated for virulence in enterohemorrhagic (Clarke *et al.*, 2006) and uropathogenic *E*. *coli* (Kostakioti *et al.*, 2009), as well as in *Salmonella enterica* Typhimurium (Moreira *et al.*, 2010, Bearson & Bearson, 2008). However, *qseB* mutants, which would be expected to mirror *qseC* mutants, do not exhibit similar defects in terms of biofilm formation (Wang *et al.*, 2011), motility (Hughes *et al.*, 2009, Guckes *et al.*, 2013, Kostakioti *et al.*, 2009), virulence (Kostakioti *et al.*, 2009), and now cell shape (Figure 5). These differences are likely due to the fact that mutants lacking QseC, but not QseB, grow misshapen and lyse. This means that (some) functions attributed to QseBC may actually be related to ancillary effects on the PG layer. To that end, QseB does not appear to regulate cell wall modifying genes (Sperandio *et al.*, 2002, Pasupuleti *et al.*, 2018, Gou *et al.*, 2019), suggesting that QseB overactivation indirectly affects cell wall synthesis. In summary, we recommend that conclusions based on *qseC* mutants be reevaluated to account for indirect effects on the cell wall, ideally by comparing *qseC* to *qseB* mutants.

### An expanding view of the RprA regulon

How does RprA overexpression reverse the shape defect of Δ*qseC* cells? While we were surprised that RprA functioned independent of its established targets to suppress the shape defect of Δ*qseC* cells, similar effects have also been observed for RprA inhibition of the CpxAR two-component signal transduction pathway (Vogt *et al.*, 2014). Specifically, *rprA* overexpression represses expression of *cpxP* and *degP*, members of the *CpxR* regulon, independent of *rpoS*, *csgD*, and *ydaM* (Vogt *et al.*, 2014). Whether RprA suppresses the shape defect of Δ*qseC* cells and inhibits the Cpx pathway by the same mechanism is unknown. However, such examples strongly suggest there are unexplored regulatory pathways that RprA controls. Indeed, RprA is predicted to bind, with various affinity, at least 200 mRNAs according to the CopraRNA database (version 2.1.2) (Raden *et al.*, 2018). Thus, dissecting the RprA regulon will likely provide important insights into the mechanisms employed by bacteria to overcome (severe) cell envelope defects. Since *rprA* expression is controlled by the Rcs signal transduction pathway (Wall *et al.*, 2018), we expect this system to play an important role in cells lacking *qseC* function.

### Morphological phenotyping informs gene function and potential therapies

The number and types of assays used to study gene function vary as widely as the pathways they represent. For some experiments though, time and cost play outsized roles in determining the extent and rigor of investigation. In such cases, identifying alternative approaches based on some phenotypic trait is critical. Historically, growth has served this function [e.g., (Typas *et al.*, 2008, Nichols *et al.*, 2011)]. However, since most mutants grow normally, more recent efforts have attempted to systematically catalog morphological phenotypes (Campos *et al.*, 2018, French *et al.*, 2017, Zahir *et al*., 2019). Here we corroborate, extend, and add to the collective morphological record 19 genes (and implicate several others) whose mutants produce shape defects in a matter of hours using relatively inexpensive reagents (i.e., LB). Moreover, the profound morphologies generated by these mutants in the Δ3PP background obviate the need for more specialized and quantitative analysis, and may serve as a useful proxy to assay the function of some of these genes.

Finally, the morphological data provided by this study may inform future, life-saving therapies. For example, increasing temperature above 37°C exacerbates the phenotype of *envZ*, *gor*, *ompR*, or *qseC* mutants (Figure 3 and S3). Since bacterial infections can induce sustained febrile responses in excess of 39°C (Ogoina, 2011), inhibitors against these proteins during febrile episodes may potentiate established treatments to shorten the duration and effects of certain bacterial infections. Thus, morphological phenotyping should be considered when designing new antibiotic treatment regimens.

## Materials and methods

### Media

Cells were cultured in LB Miller broth (1% tryptone, 0.5% yeast extract, and 1% NaCl; IBI Scientific) or λYM broth (1% tryptone, 0.1% yeast extract, 0.25% NaCl, and 0.2% maltose). Plates contained 1.5% agar (Difco). As required, antibiotics were used at the following concentrations: 100 µg ml^-1^ ampicillin, 20 µg ml^-1^ chloramphenicol, 50 µg ml^-1^ kanamycin, and 10 µg ml^-1^ tetracycline.

### Strains, plasmids, and primers

All strains, plasmids, and primers are listed in Tables S2, S3, and S4 in the supporting information, respectively. All oligonucleotides primers were obtained from Eurofins Genomics.

### Strain and plasmid construction

*E*. *coli* MG1655 Δ*lacIZYA* is the parent strain for this study (Bernhardt & de Boer, 2004). Gene deletions were constructed by using lambda Red recombination (Datsenko & Wanner, 2000) or P1-mediated transduction (Miller, 1972). Kanamycin resistance markers were evicted by using FLP recombinase produced from pCP20 (Cherepanov & Wackernagel, 1995). All gene deletions were verified by PCR. Plasmid construction is detailed in SI Text in the supporting information. All reference sequences were obtained from the Ecocyc database (Keseler *et al.*, 2017). PCR fragments were purified by using the DNA clean and concentrator kit (Zymo Research). Plasmids were purified by using the Qiaprep spin miniprep kit (Qiagen).

### Phage mutagenesis and screen to identify Δ3PP synthetic interactions

Overnight cultures of MAJ876 [Δ*lacIZYA* Δ*lpxT* Δ*ybjG* Δ*bacA*/*cat* P_lac_::*bacA lacZ*] were washed and resuspended in λYM broth. Cells were diluted 1:10 into the same medium and mutagenized with mini-Tn*10* by infecting with phage λ1098 (Way *et al.*, 1984) at a multiplicity of infection of 0.5. After 30 minutes at room temperature, cells were grown for 1.5 h at 37°C. Cells were then plated on LB agar containing tetracycline, chloramphenicol, and 500 µM IPTG (isopropyl β-D-1-thiogalactopyranoside; Research Products International) and incubated at 37°C overnight. The phage mutagenesis yielded approximately 50,000 colonies that were resuspened in LB medium containing 500 µM IPTG and frozen as described previously (Bernhardt & de Boer, 2004).

The mutant library was screened by preparing a 1 × 10^-6^ dilution in LB medium and plating on LB agar containing 60 µg ml^-1^ X-Gal (5-bromo-4-chloro-3-indolyl-β-D-galactopyranoside, Abcam) and 500 µM IPTG. [Note: tetracycline was omitted to prevent selection of insertions in pMAJ95]. Plates were incubated at 30°C, 37°C, or 42°C until X-Gal staining was clearly visible (i.e., usually 24 – 48 h). We screened between 60,000 and 150,000 colonies at each temperature, and purified those colonies that appeared solid-blue on LB agar containing 60 µg ml^-1^ X-gal and 500 µM IPTG. The positions of mini-Tn*10* insertions were determined by arbitrary PCR (O’Toole & Kolter, 1998) as described previously (Bernhardt & de Boer, 2004) using primer pairs P140/P695 (first amplification) and P141/P696 (second amplification).

### Colony phenotyping Δ3PP strain derivatives

Overnight cultures carrying pMAJ95 were diluted 1 × 10^-6^ in LB medium and plated on LB agar containing 60 µg ml-1 X-Gal and 500 µM IPTG. Plates were incubated at 30°C, 37°C, or 42°C as described above and imaged by using an iPhone X (Apple Inc.).

### Morphological phenotyping Δ3PP strain derivatives

Overnight cultures were diluted 1:2,000 in LB medium or LB medium containing chloramphenicol (to select for pMAJ95) and 500 µM IPTG and grown at 37°C or 42°C to mid-exponential phase. Live cells were then spotted on 1% agarose pads and imaged by phase-contrast microscopy by using an Olympus BX60 microscope. Images were captured by using an XM10 monochrome camera. Live cells were also prepared for flow cytometry as described previously (Jorgenson *et al.*, 2016). Morphological measurements were made using the cellSens Dimensions software (Olympus).

### Isolating multicopy suppressors of Δ*qseC*

A pBR322-based *E*. *coli* library (Ulbrandt *et al.*, 1997) was introduced into MAJ973 (Δ3PPΔ*qseC*/pMAJ95) by electroporation. Transformants were plated onto LB-IX plates containing ampicillin and 500 µM IPTG and incubated at 42°C overnight. Approximately 30,000 colonies were screened and those colonies that appeared light blue, indicating loss of pMAJ95, were purified. From these, we observed the formation of small- and large-colony variants. Overnight cultures of cells from the small and large-colony variants were diluted 1:2,000 in LB medium containing ampicillin and 500 µM IPTG, grown at 42°C to an optical density at 600 nm (OD_600_) of ∼0.5, and imaged under phase-contrast microscopy. Microscopic analysis revealed that large colony variants produced rod-shaped cells whereas small colony variants produced misshapen cells. Plasmids were isolated from the large-colony variants, transformed into MAJ1005 (Δ*qseC*), and tested for their ability to reverse the shape defect of Δ*qseC* cells as above. Suppressing plasmids were sequenced using primers P735 and P736.

## Acknowledgements

We thank Susan Gottesman and Nadim Majdalani (NCI) for providing the pORF library, plasmids pBRplac and p*rprA*, and for technical guidance. We thank Kevin Young for helpful discussions during the preparation of this manuscript. We also thank Anne Darley and Victoria Marcelle for technical assistance. This study was supported by the National Institute of General Medical Sciences NIH grant GM061019. The UAMS DNA core facility is supported in part by the Center for Microbial Pathogenesis and Host Inflammatory Responses NIH grant GM103625. The content is solely the responsibility of the authors and does not necessarily represent the official views of the NIH. The authors have no conflicts of interest to declare.

