## Supporting Information for "A genetic screen to identify factors affected by undecaprenyl phosphate recycling uncovers novel connections to morphogenesis in *Escherichia coli*"

**<u>Supplemental information for:</u>**

*Corresponding author

**Running title**: Und-P synthetic interactions

**Key words**: BacA, bactoprenol, lysis, morphology, peptidoglycan, QseC

### Supplemental methods

**Plasmid construction.** pMAJ95 ($P_{lac}$::*bacA*) is a plasmid that expresses *bacA* from the unstable plasmid, and was constructed by amplifying 75 bps upstream and 50 bps downstream of the *bacA* open reading frame with primers P545 and P546. The 963 bp product was cut with EcoRI and HindIII and ligated into the same sites of pTB238 (Paradis-Bleau *et al.*, 2010). pMAJ117 ($P_{204}$::*ompR*) is a plasmid that expresses *ompR*, and was constructed by amplifying the *ompR* open reading frame with primers P662 and P663. The 735 bp product was cut with EcoRI and PstI and ligated to the same sites of pDSW361. pMAJ123 ($P_{204}$::*qseC*) is a plasmid that expresses *qseC*, and was constructed by amplifying the *qseC* open reading frame with primers P737 and P738. The 1,365 bp product was cut with EcoRI and PstI and ligated to the same site of pDSW361. pMAJ131 ($P_{204}$::*gor*) is a plasmid that expresses *gor*, and was constructed by amplifying the *gor* open reading frame with primers P864 and P865. The 1,368 bp product was cut with EcoRI and PstI and ligated to the same sites of pDSW361. pMAJ133 ($P_{204}$::*pdhR*) is plasmid that expresses *pdhR*, and was constructed by amplifying the *pdhR* open reading frame with primers P940 and P941. The 780 bp product was cut with EcoRI and PstI and ligated to the same sites of pDSW361. pMAJ139 ($P_{lac}$::*qseC*) is a plasmid that expresses *qseC*, and was constructed by amplifying the *qseC* open reading frame with primers P957 and P962. The 1,368 bp product was cut with EcoRI and HindIII and ligated to the same sites of pBRplac. pMAJ140 ($P_{204}$::*envZ*) is a plasmid that expresses *envZ*, and was constructed by amplifying the *envZ* open reading frame with primers P967 and P968. The 1,368 bp product was cut with SacI and HindIII and ligated to the same sites of pDSW361. All plasmid constructs were verified by DNA sequencing at the UAMS DNA Sequencing Core Facility.

**Table S1. Mini-Tn*10* location in Δ3PP mutants**

| Gene | Location |
|---|---|
| *ftsK* | N1051: AA*C |
| *lon* | V355*G356 |
| *minD* | M196: A*TG |
| *minE* | 38 bps after *minE* stop codon |
| *mreB* | 261 bps upstream of *mreB* start codon |
| *seqA* | R73: CG*T |
| *topA* | K727*Y728 |

*Denotes location of mini-Tn*10*

**Table S2. Strains used in this study**

| Strain | Relevant features | Source or reference |
|---|---|---|
| MAJ25 | MG1655 *lacIZYA*::*frt*; strain is TB28 | (Bernhardt & de Boer, 2004) |
| MAJ815 | MAJ25 Δ*lpxT*::*frt* Δ*ybjG*::*frt* Δ*bacA*::*frt* | This study |
| MAJ876 | MAJ815/pMAJ95 | This study |
| MAJ890 | MAJ876 Δ*ompR*::*kan* | This study |
| MAJ891 | MAJ876 Δ*envZ*::*kan* | This study |
| MAJ916 | MAJ876 Δ*gor*::*kan* | This study |
| MAJ936 | MAJ25 $P_{mreB}$::mini-Tn*10* | This study |
| MAJ937 | MAJ876 $P_{mreB}$::mini-Tn*10* | This study |
| MAJ938 | MAJ25 Δ*ompR*::*frt* | This study |
| MAJ941 | MAJ938/pDSW361 | This study |
| MAJ962 | MAJ25 Δ*qseB*::*cat* | This study |
| MAJ964 | MAJ25 Δ*envZ*::*kan* | This study |
| MAJ969 | MAJ25 Δ*gor*::*kan* | This study |
| MAJ973 | MAJ876 Δ*qseC*::*kan* | This study |
| MAJ974 | MAJ876 Δ*pgpB*::*kan* | This study |
| MAJ983 | MAJ876 Δ*mrcB*::*kan* | This study |
| MAJ999 | MAJ25 Δ*qseBC*::*kan* | This study |
| MAJ1002 | MAJ25 Δ*pmrB*::*frt* | This study |
| MAJ1003 | MAJ1002 Δ*qseC*::*kan* | This study |
| MAJ1005 | MAJ25 Δ*qseC*::*frt* | This study |
| MAJ1015 | MAJ1005/pMAJ45 | This study |
| MAJ1016 | MAJ1005/pDSW361 | This study |
| MAJ1021 | MAJ1005/pMAJ123 | This study |
| MAJ1022 | MAJ25 Δ*pmrA*::*cat* | This study |
| MAJ1023 | MAJ1005 Δ*pmrA*::*cat* | This study |
| MAJ1091 | MAJ876 Δ*wzxE*::*kan* | This study |
| MAJ1092 | MAJ876 Δ*cpxR*::*kan* | This study |
| MAJ1093 | MAJ876 Δ*cpxR*::*kan* | This study |
| MAJ1094 | MAJ876 Δ*pal*::*kan* | This study |
| MAJ1095 | MAJ876 Δ*mltG*::*kan* | This study |
| MAJ1096 | MAJ876 Δ*rep*::*kan* | This study |
| MAJ1097 | MAJ876 Δ*ldcA*::*kan* | This study |
| MAJ1110 | MAJ876 Δ*lpoB*::*kan* | This study |
| MAJ1115 | MAJ876 Δ*dacA*::*kan* | This study |
| MAJ1125 | MAJ876 *ftsK*::mini-Tn*10* | This study |
| MAJ1126 | MAJ876 *topA*::mini-Tn*10* | This study |
| MAJ1136 | MAJ876 Δ*seqA*::*kan* | This study |
| MAJ1140 | MAJ25 Δ*gor*::*frt*/pMAJ131 | This study |
| MAJ1142 | MAJ25 Δ*gor*::*frt*/pDSW361 | This study |
| MAJ1143 | MAJ1005/pBRplac | This study |
| MAJ1144 | MAJ1005/pRprA | This study |

| | | |
|---|---|---|
| MAJ1146 | MAJ876 *lon*::mini-Tn*10* | This study |
| MAJ1147 | MAJ876 $T_{minE}$::mini-Tn*10* | This study |
| MAJ1148 | MAJ876 *minD*::mini-Tn*10* | This study |
| MAJ1151 | MAJ1005 Δ*rpoS*::*kan*/pRprA | This study |
| MAJ1155 | MAJ1005/pMAJ133 | This study |
| MAJ1163 | MAJ1005 Δ*hfq*::*kan*/pRprA | This study |
| MAJ1170 | MAJ25 Δ*hfq*::*kan* | This study |
| MAJ1176 | MAJ938/pMAJ117 | This study |
| MAJ1177 | MAJ1005/pMAJ139 | This study |
| MAJ1188 | MAJ25 Δ*envZ*::*frt*/pMAJ140 | This study |
| MAJ1189 | MAJ25 Δ*envZ*::*frt*/pDSW361 | This study |
| MAJ1197 | MAJ1005 Δ*ydaM*::*kan*/pRprA | This study |
| MAJ1198 | MAJ1005 Δ*csgD*::*kan*/pRprA | This study |
| MAJ1239 | MAJ25 Δ*ygiW*::*kan* | This study |
| MAJ1261 | MAJ25 Δ*ygiW*::*frt* Δ*qseC*::*kan* | This study |
| **Additional *E. coli* strains used as donors for P1 mediated transduction** | | |
| AV21-10K | MG1655 Δ*dacA*::*kan* | (Varma & Young, 2004) |
| JW0674 | BW25113 Δ*seqA*::*kan* | (Baba *et al.*, 2006) |
| JW0731 | BW25113 Δ*pal*::*kan* | (Baba *et al.*, 2006) |
| JW1023 | BW25113 Δ*csgD*::*kan* | (Baba *et al.*, 2006) |
| JW1083 | BW25133 Δ*mltG*::*kan* | (Baba *et al.*, 2006) |
| JW1181 | BW25113 Δ*ldcA*::*kan* | (Baba *et al.*, 2006) |
| JW2992 | BW25113 Δ*ygiW*::*kan* | (Baba *et al.*, 2006) |
| JW3367 | BW25113 Δ*envZ*::*kan* | (Baba *et al.*, 2006) |
| JW3368 | BW25113 Δ*ompR*::*kan* | (Baba *et al.*, 2006) |
| JW3883 | BW25113 Δ*cpxR*::kan | (Baba *et al.*, 2006) |
| JW4130 | BW25113 Δ*hfq*::*kan* | (Baba *et al.*, 2006) |
| JW5206 | BW25113 Δ*ydaM*::*kan* | (Baba *et al.*, 2006) |
| JW5604 | BW25113 Δ*rep*::*kan* | (Baba *et al.*, 2006) |
| MAJ97 | CS109 Δ*wzxE*::*kan* | (Jorgenson *et al.*, 2016) |
| MAJ657 | MG1655 Δ*mrcB*::*kan* | (Jorgenson *et al.*, 2019) |
| MAJ659 | MG1655 Δ*lpoB*::*kan* | (Jorgenson *et al.*, 2019) |
| MAJ907 | MG1655 Δ*pgpB*::*kan* | This study |
| MAJ908 | MG1655 Δ*gor*::*kan* | This study |
| MAJ963 | MAJ25 Δ*qseC*::*kan* | This study |

**Table S3. Plasmids used in this study**

| Plasmid | Relevant genotype or characteristics | Origin of replication | Source or reference |
|---|---|---|---|
| pCP20 | $\lambda_{PR}$::*flp* λcI857 *bla cat* Rep(Ts) | pSC101 | (Cherepanov & Wackernagel, 1995) |
| pBRplac | $P_{lac}$ $Amp^R$ $Tet^R$ | pBR | (Guillier & Gottesman, 2006) |
| pDSW361 | $P_{204}$ *lacI*$^q$ *kan* $Kan^R$ derivative of pDSW204. | pBR | (Weiss *et al.*, 1999) |
| pKD3 | *bla frt-cat-frt* | R6Kγ | (Datsenko & Wanner, 2000) |
| pKD13 | *bla frt-aph-frt* | R6Kγ | (Datsenko & Wanner, 2000) |
| pKD46 | $P_{araB}$::*gam-bet-exo bla* Rep(Ts) | pSC101 | (Datsenko & Wanner, 2000) |
| pMAJ45 | $P_{204}$::*uppS* | pBR | (Jorgenson & Young, 2016) |
| pMAJ95 | $P_{lac}$::*bacA lacZYA repE cat* | oriS | This study |
| pMAJ117[a] | $P_{204}$::*ompR* | pBR | This study |
| pMAJ123[a] | $P_{204}$::*qseC* | pBR | This study |
| pMAJ131 | $P_{204}$::*gor* | pBR | This study |
| pMAJ133 | $P_{204}$::*pdhR* | pBR | This study |
| pMAJ139[b] | $P_{lac}$::*qseC* | pBR | This study |
| pMAJ140[a] | $P_{204}$::*envZ* | pBR | This study |
| pRprA[b] | $P_{lac}$::*rprA* | pBR | (Soper *et al.*, 2010) |

[a]Derivative of pDSW361.

[b]Derivative of pBRplac.

**Table S4. Primers used in this study**

| Primer | Sequence[a] | Purpose |
|---|---|---|
| P122 | ACGCAGTAGTTCGGACAAGCGGTACATTTTAATAATTTAGAT TCCGGGGATCCGTCGACC | Δ*bacA* |
| P123 | CAATCGTTGACAACGCCAAGCATCCGACACTATTCCTCAAT GTAGGCTGGAGCTGCTTCG | Δ*bacA* |
| P140 | GTTGTGATGTAGGCATCAATC | Mapping |
| P141 | AAGGCACCTTTGGTCACCAAC | Mapping |
| P533 | GTTGCCTGCGTTTTTCAGTAAGATAATTAGAGAAAATATGAT TCCGGGGATCCGTCGACC | Δ*lpxT* |
| P534 | TGATGTTAATTACTGTGAGTTATTTGTTTTGGAAATGTTTTGT AGGCTGGAGCTGCTTCG | Δ*lpxT* |
| P537 | TATTGGCTCCCTTTTAATCACTTTGCGTCGGGAAGTTATGTG TAGGCTGGAGCTGCTTCG | Δ*ybjG* |
| P538 | CTACACTTTTTAGCAGAGATCAGTCACGCACCCAGCCTTTAT TCCGGGGATCCGTCGACC | Δ*ybjG* |
| P545 | CAGGAATTCCCAAACGGTTATAACCTGGTC | pMAJ95 |
| P546 | TTGAAGCTTTCGGCCTACGCAATCGTTGAC | pMAJ95 |
| P650 | CAGTTATGATTGTGGGACTTATCAAAAAGGAGAGGCCATGA TTCCGGGGATCCGTCGACC | Δ*pgpB* |
| P651 | TTCGGAAAATCAACCAGCGTTAACTTTCTTGTTCTCGTTGTG TAGGCTGGAGCTGCTTCG | Δ*pgpB* |
| P654 | CTACAATCGCGGTAATCAACGATAAGGACACTTTGTCATGT GTAGGCTGGAGCTGCTTCG | Δ*gor* |
| P655 | ACTCTTAGCCCTTTAACATTTAACGCATTGTCACGAACTCAT TCCGGGGATCCGTCGACC | Δ*gor* |
| P662 | CAAGAATTCCAAGAGAACTACAAGATTCTG | pMAJ117 |
| P663 | TTGCTGCAGTCATGCTTTAGAGCCGTCCGG | pMAJ117 |
| P695 | GGCCACGCGTCGACTAGTACNNNNNNNNNNGATGC | Mapping |
| P696 | GGCCACGCGTCGACTAGTAC | Mapping |
| P714 | TACTGGCAGCAAATACGGTTATCGCAGGGATGAAAAAATGT GTAGGCTGGAGCTGCTTCG | Δ*qseB* |
| P715 | GCGTCAGCCTGACGCGCAGACTAAGACGTTGGGTAAATTTC ATATGAATATCCTCCTTAG | Δ*qseB* |
| P718 | CCGTGCATGGTATTGGTTACACATTAGGTGAGAAATGAAAA TTCCGGGGATCCGTCGACC | Δ*qseC* |
| P719 | GCTCTCATAGACAGAGAAGTTACCAGCTTACCTTCGCCTCT GTAGGCTGGAGCTGCTTCG | Δ*qseC* |
| P726 | TATGCTGGTCGCGAATGAGGAAAACTAATTGAATCTGATGT GTAGGCTGGAGCTGCTTCG | Δ*pmrB* |
| P727 | TGCCGGAAGTCTCTTGCCGGTTTTGCAGGAAAAACTGCCCA TTCCGGGGATCCGTCGACC | Δ*pmrB* |
| P735 | CCTGACGTCTAAGAAACCATTATTATC | pBR322 |
| P736 | AACGACAGGAGCACGATCATGCG | pBR322 |
| P737 | CAGGAATTCAAATTTACCCAACGTCTTAGTC | pMAJ123 |

| | | |
|---|---|---|
| P738 | TTGCTGCAGTTACCAGCTTACCTTCGCCTC | pMAJ123 |
| P741 | CGGATGATATTCTGCAAACTTGCAGGAGAGTGAGTGAATGT GTAGGCTGGAGCTGCTTCG | Δ*pmrA* |
| P742 | TCAGCCGTTGGCGCAGCGATATTGGTCGGCGCAGAAAATG CATATGAATATCCTCCTTAG | Δ*pmrA* |
| P864 | CAGGAATTCACTAAACACTATGATTACATC | pMAJ131 |
| P865 | TTGCTGCAGTTAACGCATTGTCACGAACTC | pMAJ131 |
| P940 | CAAGAATTCGCCTACAGCAAAATCCGCCAAC | pMAJ133 |
| P941 | CTGCTGCAGCTAATTCTTTCGTTGCTCCAG | pMAJ133 |
| P957 | TTGAAGCTTTTACCAGCTTACCTTCGCCTC | pMAJ139 |
| P962 | CAGGAATTCATGAAATTTACCCAACGTCTTAG | pMAJ139 |
| P967 | CAGGAGCTCAGGCGATTGCGCTTCTCGCCAC | pMAJ140 |
| P968 | TTGCTGCAGTTACCCTTCTTTTGTCGTGCC | pMAJ140 |

[a]All primer sequences are written 5'→ 3'. Restriction sites are underlined.

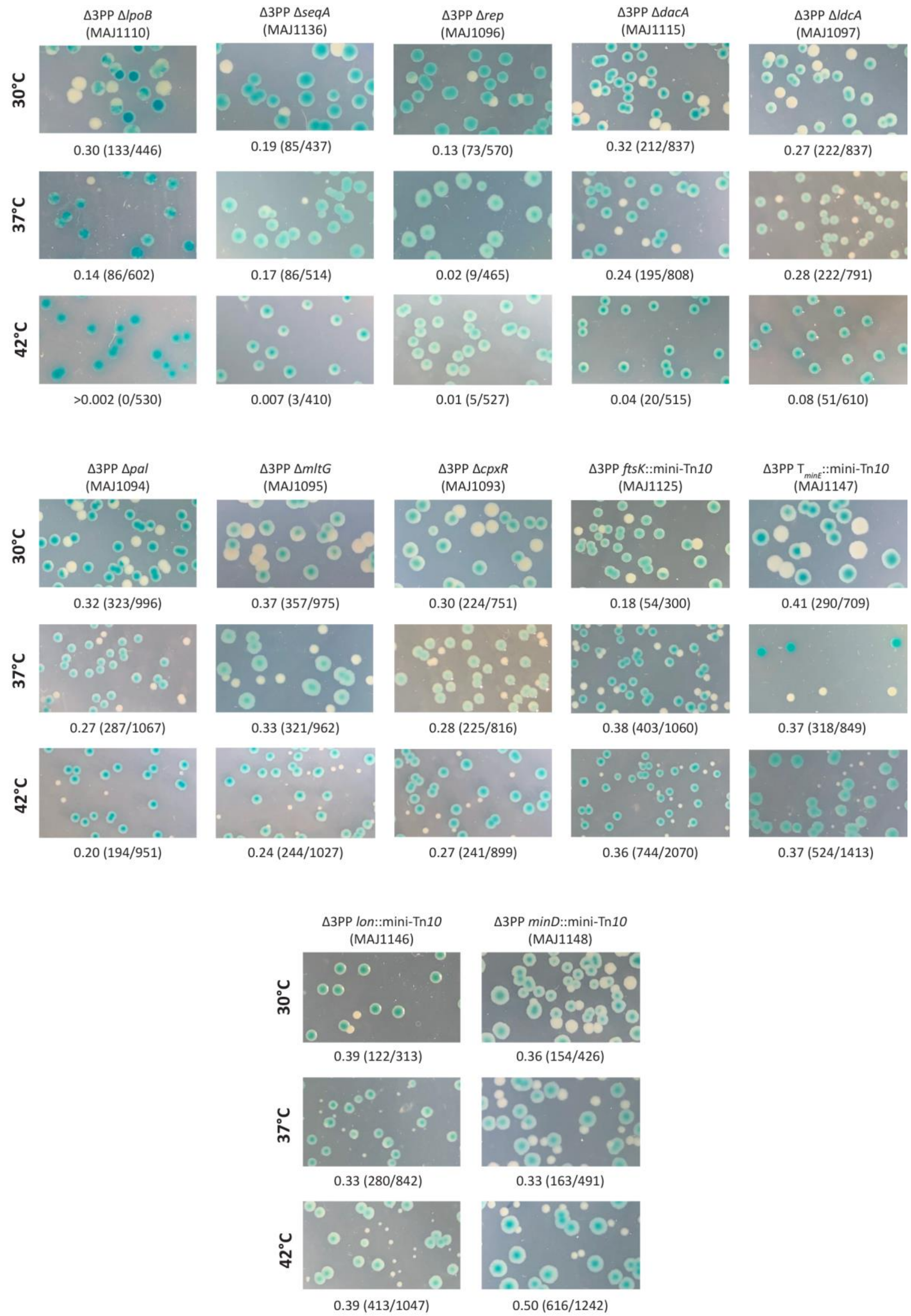

Δ3PP ΔlpoB (MAJ1110)
Δ3PP ΔseqA (MAJ1136)
Δ3PP Δrep (MAJ1096)
Δ3PP ΔdacA (MAJ1115)
Δ3PP ΔldcA (MAJ1097)
30°C
0.30 (133/446)
0.19 (85/437)
0.13 (73/570)
0.32 (212/837)
0.27 (222/837)
37°C
0.14 (86/602)
0.17 (86/514)
0.02 (9/465)
0.24 (195/808)
0.28 (222/791)
42°C
>0.002 (0/530)
0.007 (3/410)
0.01 (5/527)
0.04 (20/515)
0.08 (51/610)
Δ3PP Δpal (MAJ1094)
Δ3PP ΔmltG (MAJ1095)
Δ3PP ΔcpxR (MAJ1093)
Δ3PP ftsK::mini-Tn10 (MAJ1125)
Δ3PP $T_{minE}$::mini-Tn10 (MAJ1147)
30°C
0.32 (323/996)
0.37 (357/975)
0.30 (224/751)
0.18 (54/300)
0.41 (290/709)
37°C
0.27 (287/1067)
0.33 (321/962)
0.28 (225/816)
0.38 (403/1060)
0.37 (318/849)
42°C
0.20 (194/951)
0.24 (244/1027)
0.27 (241/899)
0.36 (744/2070)
0.37 (524/1413)
Δ3PP lon::mini-Tn10 (MAJ1146)
Δ3PP minD::mini-Tn10 (MAJ1148)
30°C
0.39 (122/313)
0.36 (154/426)
37°C
0.33 (280/842)
0.33 (163/491)
42°C
0.39 (413/1047)
0.50 (616/1242)

**Figure S1. Viability of (additional) Δ3PP mutant derivatives.** Effect of various mutations on the segregation of pMAJ95 in Δ3PP cells. Colonies appear solid blue in mutants that are synthetically lethal in the Δ3PP background (i.e., Δ3PP Δ*lpoB* 42°C) because plasmid-free cells lyse. The frequency of pMAJ95 segregation is reduced in mutants that are synthetically sick in the Δ3PP background because pMAJ95 confers a growth advantage. Above photographs: genotype and strain designation. Below photographs: fraction of white colonies to the total number of colonies.

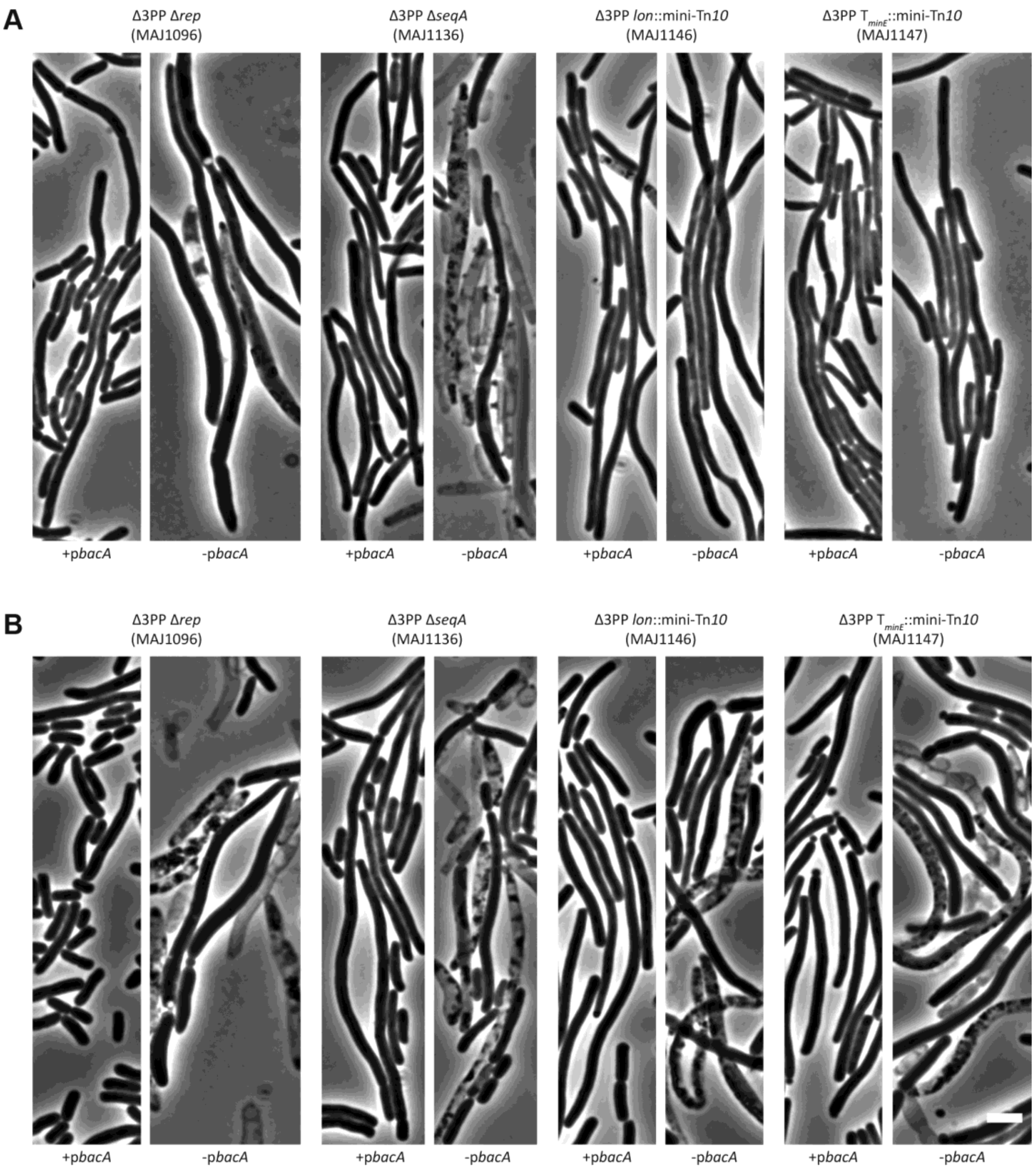


**Figure S2. Morphology of Δ3PP mutant filaments.** Cells with the indicated genotypes were grown at 37°C (A) or 42°C (B) in LB (-p*bacA*) or LB containing chloramphenicol and 500 µM IPTG (+p*bacA*) until the culture reached an $OD_{600}$ of 0.3-0.4. The cells were then photographed by phase-contrast microscopy. The white bar represents 3 µm. The data is representative of at least two independent experiments. Above photographs: genotype and strain designation.

111

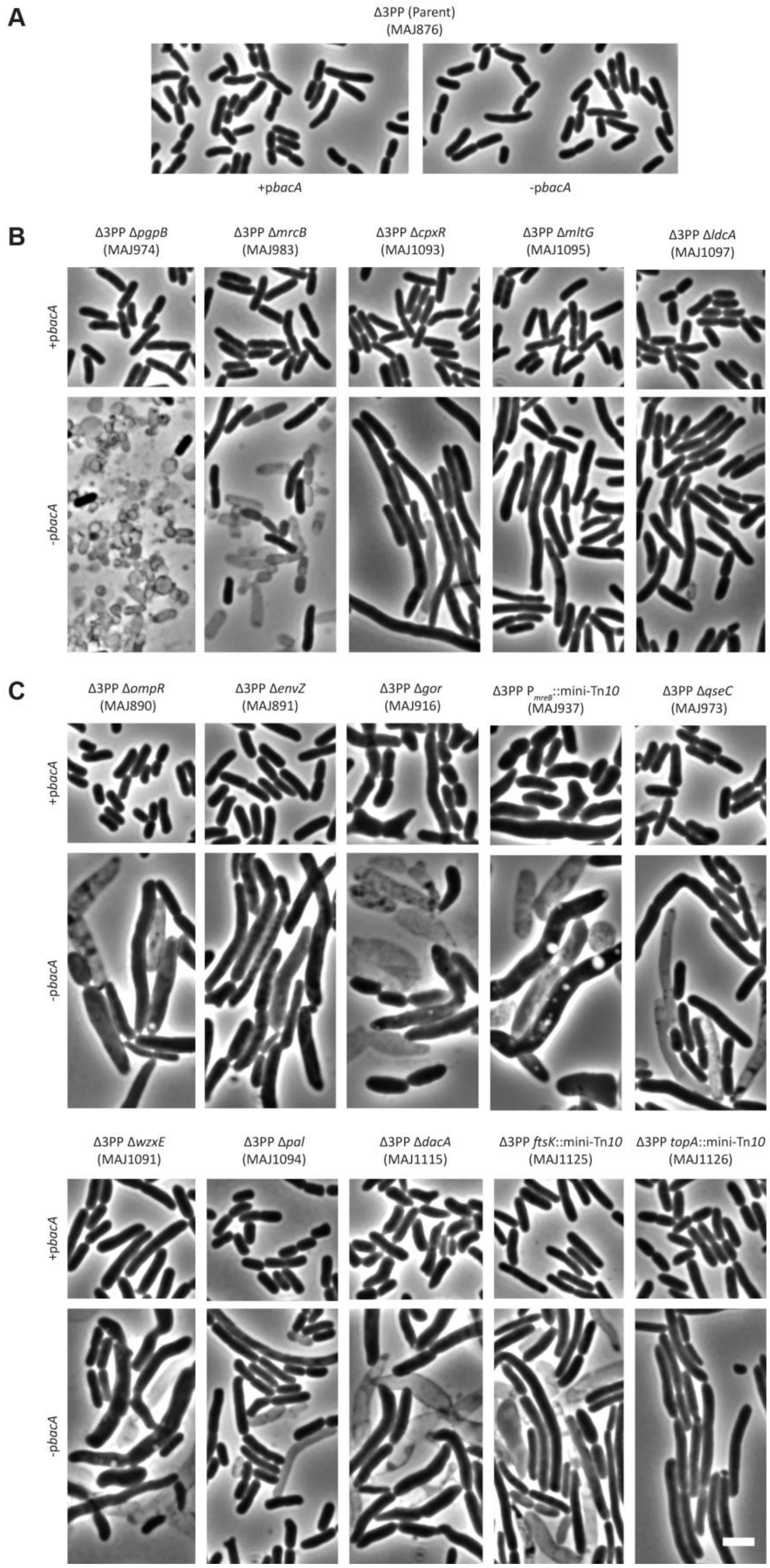
A
Δ3PP (Parent)
(MAJ876)
+pbacA
-pbacA
B
Δ3PP ΔpgpB
(MAJ974)
Δ3PP ΔmrcB
(MAJ983)
Δ3PP ΔcpxR
(MAJ1093)
Δ3PP ΔmltG
(MAJ1095)
Δ3PP ΔldcA
(MAJ1097)
+pbacA
-pbacA
C
Δ3PP ΔompR
(MAJ890)
Δ3PP ΔenvZ
(MAJ891)
Δ3PP Δgor
(MAJ916)
Δ3PP P mreB::mini-Tn10
(MAJ937)
Δ3PP ΔqseC
(MAJ973)
+pbacA
-pbacA
Δ3PP ΔwzxE
(MAJ1091)
Δ3PP Δpal
(MAJ1094)
Δ3PP ΔdacA
(MAJ1115)
Δ3PP ftsK::mini-Tn10
(MAJ1125)
Δ3PP topA::mini-Tn10
(MAJ1126)
+pbacA
-pbacA

**Figure S3. Morphology of Δ3PP mutant derivatives at 37°C.** Cells with the indicated genotypes were grown at 37°C in LB (-p*bacA*) or LB containing chloramphenicol and 500 µM IPTG (+p*bacA*) until the culture reached an $OD_{600}$ of 0.3-0.4. The cells were then photographed by phase-contrast microscopy. The white bar represents 3 µm. (A) The Δ3PP parent. (B) Mutant derivatives that produce shape defects in the absence of *bacA* expression. (C) Mutant derivatives that produce shape defects independent of *bacA* expression. The data is representative of at least two independent experiments. Above photographs: genotype and strain designation.

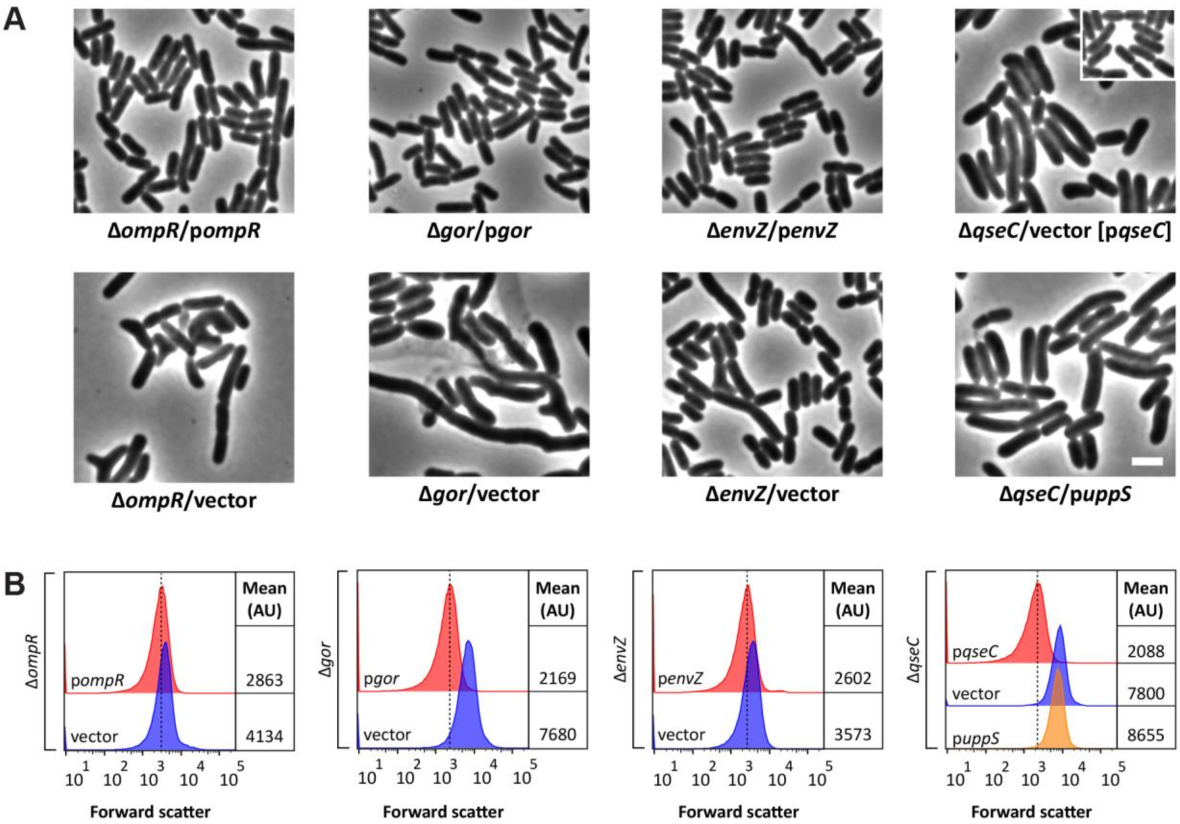


**Figure S4. Complementation of *envZ*, *gor*, *ompR*, and *qseC* mutants.** (A) Micrographs of Δ*ompR*, Δ*gor*, Δ*envZ*, or Δ*qseC* cells containing derivatives of pDSW361 that express the indicated genes. Cells were grown at 37°C in LB until the culture reached an $OD_{600}$ of 0.5. The cells were then photographed by phase-contrast microscopy. [p*qseC*]: inset showing the effect of overexpressing *qseC* in Δ*qseC* cells. The white bar represents 3 µm. (B) Live cells from panel A were also examined by flow cytometry. Histograms of the forward scatter area from 100,000 events (cells) are shown. The mean cell size of the wild type is represented by the dashed line and is expressed in arbitrary units (AU). The strains shown are MAJ1176 (Δ*ompR*/p*ompR*), MAJ941 (Δ*ompR*/vector), MAJ1140 (Δ*gor*/p*gor*), MAJ1142 (Δ*gor*/vector), MAJ1188

(Δ*envZ*/p*envZ*), MAJ1189 (Δ*envZ*/vector), MAJ1021 (Δ*qseC*/p*qseC*), MAJ1016 (Δ*qseC*/vector), and MAJ1015 (Δ*qseC*/p*uppS*).

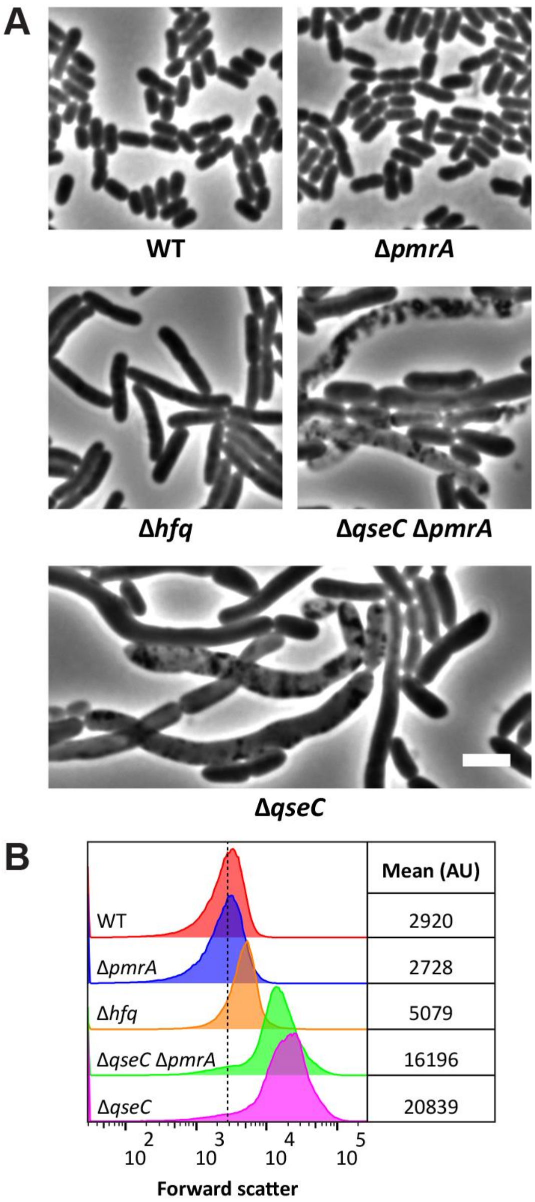


**Figure S5. Morphology of *pmrA* and *hfq* mutants.** (A) Cells with the indicated genotypes were grown at 42°C in LB until the culture reached an $OD_{600}$ of 0.5. The cells were then photographed by phase-contrast microscopy. The white bar represents 3 µm. (B) Live cells from panel A were also examined by flow cytometry. Histograms of the forward scatter area from 100,000 events (cells) are shown. The mean cell size of the wild type is represented by the dashed line and is expressed in arbitrary units (AU).

Data are representative of two independent experiments.  The strains shown are MAJ25 (WT), MAJ1022 (Δ*pmrA*), MAJ1170 (Δ*hfq*), MAJ1023 (Δ*qseC* Δ*pmrA*), and MAJ1005 (Δ*qseC*)

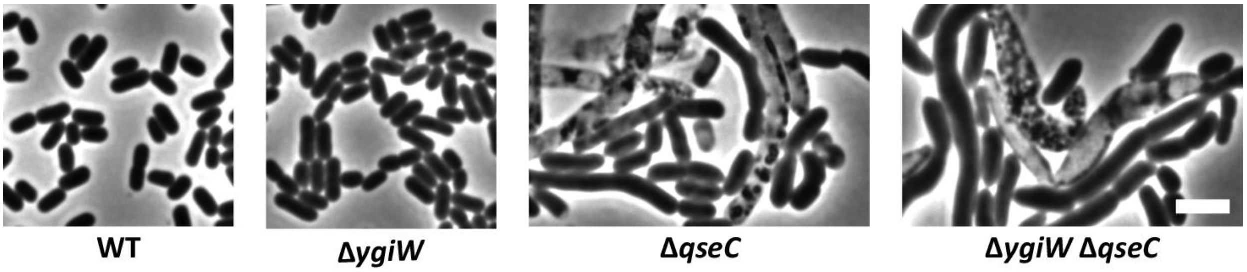


**Figure S6. Deleting *ygiW* does not reverse the shape defect of Δ*qseC* cells.** Cells with the indicated genotypes were grown at 42°C in LB until the culture reached an $OD_{600}$ of 0.5. The cells were then photographed by phase-contrast microscopy. The white bar represents 3 µm. The strains shown are MAJ25 (WT), MAJ1239 (Δ*ygiW*), MAJ1005 (Δ*qseC*), and MAJ1261 (Δ*ygiW* Δ*qseC*).

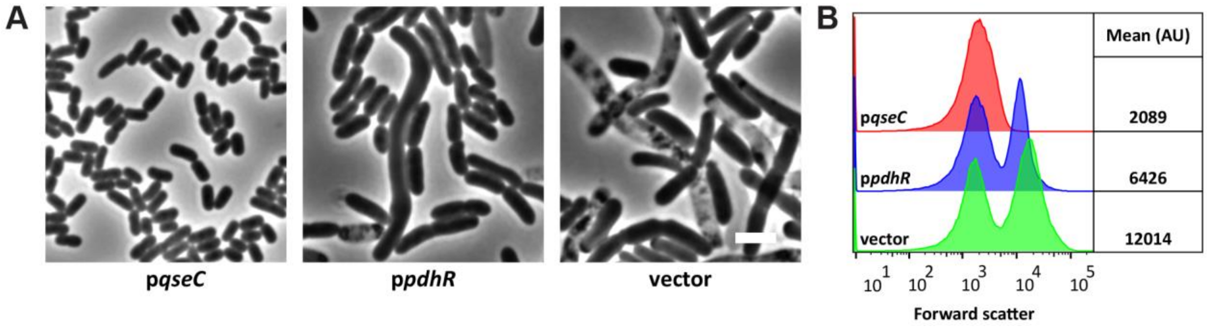


**Figure S7. Multicopy *pdhR* partially suppresses the shape defect of Δ*qseC* cells.** (A) Micrographs of Δ*qseC* cells containing derivatives of pDSW361 that express *qseC* or *pdhR*. Cells were grown at 42°C in LB until the culture reached an $OD_{600}$ of 0.5. The cells were then photographed by phase-contrast microscopy. The white bar represents 3 µm. (B) Live cells from panel A were also examined by flow cytometry. Histograms of the forward scatter area from 100,000 events (cells) are shown. The mean cell size of the wild type is represented by the dashed line and is expressed in arbitrary units (AU). Data are representative of two independent experiments. The strains shown are MAJ1021 (p*qseC*), MAJ1155 (p*pdhR*), and MAJ1016 (vector).

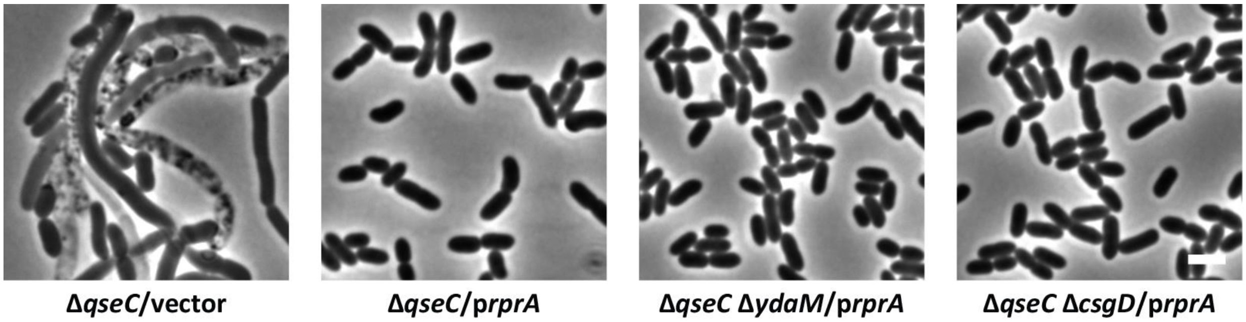


**Figure S8. Multicopy *rprA* does not require YdaM or CsgD to suppress the shape defect of Δ*qseC* cells.** Micrographs of cells with the indicated genotypes containing pBRplac (vector) or p*rprA*. Cells were grown at 42°C in LB until the culture reached an $OD_{600}$ of 0.5. The cells were then photographed by phase-contrast microscopy. The white bar represents 3 µm. The strains shown are MAJ1143 (Δ*qseC*/vector), MAJ1144 (Δ*qseC/*p*rprA*), MAJ1197 (Δ*qseC* Δ*ydaM*/p*rprA*), and MAJ1198 (Δ*qseC* Δ*csgD*/p*rprA*).
